# Sequential recruitment of accumbal dopamine and acetylcholine during learning

**DOI:** 10.1101/2025.01.06.631529

**Authors:** Zhewei Zhang, Kauê Machado Costa, Wenhui Zong, Yizhou Zhuo, Guochuan Li, Yulong Li, Geoffrey Schoenbaum

**Affiliations:** National Institute on Drug Abuse Intramural Research Program, National Institutes of Health, Baltimore, MD, 21224, USA; Department of Psychology, University of Alabama at Birmingham, Birmingham, AL, 35223, USA; State Key Laboratory of Membrane Biology, School of Life Sciences, Peking University, Beijing 100871, China; PKU-IDG/McGovern Institute for Brain Research, Beijing, 100871, China; Peking-Tsinghua Center for Life Sciences, New Cornerstone Science Laboratory, Academy for Advanced Interdisciplinary Studies, Peking University, Beijing, 100871, China; Chinese Institute for Brain Research, Beijing, 102206, China; Institute of Molecular Physiology, Shenzhen Bay Laboratory, Shenzhen, Guangdong, 518055, China; National Biomedical Imaging Center, Peking University, Beijing, 100871, China

**Author notes:** Equal contribution / co-first authors (Zhewei Zhang, Kauê Machado Costa).

## Abstract

Associative learning depends on neuromodulator release in the nucleus accumbens, including both dopamine and acetylcholine. Dopamine signals convey prediction errors that drive learning and dynamically evolve as predictions are acquired, but the role of acetylcholine in learning is far less clear. We hypothesized that acetylcholine signals change over the course of learning, complementing dopaminergic encoding of prediction errors. To test this hypothesis, we examined how acetylcholine and dopamine dynamics co-evolve during learning by recording these two signals as rats learned three cueguided operant tasks. We discovered that neuromodulatory responses to task cues evolve according to a specific sequential structure and correlate with different features of associative learning. When rats learn a cue-outcome association, there is first the appearance of indiscriminate cholinergic dips, then dopamine responses differentiate based on predicted value, followed by cholinergic dips also segregating according to value. Stable performance required this full pattern, and acetylcholine dips could be decoupled from value by changing reward contingencies. Moreover, optogenetic manipulation of accumbal cholinergic interneurons during cue presentation bidirectionally altered learning rate, with activation slowing and inhibition facilitating associative learning. Using computational modeling, we demonstrate that the observed acetylcholine dynamics best fit an associative salience term predicted by hybrid attentional associative learning models, suggesting that dopamine and acetylcholine act complementarily to determine the content and rate of learning.

---

Learning is critical for survival. Determining which cues predict relevant outcomes, like the presence of food or the approach of predators, is paramount for every animal. The formation of associations between cues and motivationally-relevant outcomes is driven by reward prediction errors (RPEs). Dopamine (DA) release in the nucleus accumbens (NAcc) has long been regarded as a neural instantiation of these prediction error signals ^1-4^. However, dopamine is only one of several neuromodulators that are released in the NAcc. Another important neuromodulator is acetylcholine (ACh), which in the NAcc is released primarily by local cholinergic interneurons ^5-7^. There is evidence that in the NAcc, acetylcholine regulates dopamine release and synaptic plasticity to influence learning ^8-11^, but its precise functional role is unclear.

Unlike dopamine neurons, which increase their firing in response to rewards and reward-predictive cues, most striatal cholinergic neurons are tonically active and pause their firing in response to these events^12,13^. We recently demonstrated that acetylcholine release in the NAcc core also dips in response to rewards and reward-predictive cues^14^. Similar patterns have been observed in the context of drug reward and in other regions of the striatum^15-18^, and inhibition of cholinergic interneurons in the NAcc augments cue-motivated behavior^19,20^. While there is some conflicting evidence^9,21^, the majority of the literature indicates that acetylcholine typically dips during the periods where dopamine signals reflect RPEs ^14,16,18,22^. Moreover, at the cellular level, acetylcholine typically acts in opposition to dopamine in its effects on synaptic plasticity^5,6^. Thus, these dips are well positioned to gate or modulate dopamine-induced plasticity.

Most studies on the interaction between dopamine and acetylcholine have only used causal manipulations or focused on acetylcholine signals when associations are already well learned. As a result, little is known about how acetylcholine dynamics evolve during learning, or how they align with dopamine signals, which are known to change according to the update of predictive relationships between cues and outcomes. This is critical for understanding the cognitive function of acetylcholine, as the dynamics of acetylcholine may differ depending on learning stage, and such distinctions could be obscured if observations are limited to well-trained subjects.

To uncover the role of the accumbal acetylcholine during learning, we used multi-color fiber photometry to simultaneously record acetylcholine and dopamine signals in the NAcc of rats as they learned three different odor discrimination tasks. We found that cue-evoked acetylcholine and dopamine signals evolve according to a characteristic pattern, with non-specific dips in acetylcholine preceding, and cue-specific dips following, the signaling of dopamine prediction errors. These changes in dopamine and acetylcholine signals occurred before rats reached task proficiency, suggesting that this dynamic neuromodulatory progression pattern could underlie learning. Consistent with this hypothesis, acetylcholine dips were correlated with the changes in dopamine signals and behavioral learning progression, predicting the rate of associative learning. Computational modeling showed that acetylcholine dynamics were best explained as an associative salience signal, as predicted by hybrid attentional associative learning models ^23^ that integrate principles from the Mackintosh ^24,25^ and Pearce-Hall ^26,27^ models. Predictive cues acquire this associative salience through learned associations with reward or punishment, distinct from intrinsic salience arising from inherent physical properties, such as intensity or novelty. Accordingly, optogenetic manipulation of accumbal cholinergic interneuron activity confirmed that augmenting and suppressing cholinergic dips increased and decreased the rate of associative learning, respectively. Together, these results reveal that associative learning is determined by a sequential dynamic pattern of acetylcholine and dopamine in the NAcc, where acetylcholine regulates learning by signaling the acquired associative salience of cues.

## Results

To investigate the functional role of acetylcholine in the NAcc core during the progression of associative learning, we recorded acetylcholine signals using the genetically-encoded green fluorescent sensor gACh4h^14^ (Extended Data Fig. 1A). To take advantage of the link between NAcc dopamine and RPE representation, we also simultaneously recorded dopamine signals with the red fluorescent sensor rDA3m^28^ (Extended Data Fig. 1A). These sensors were monitored in the NAcc of freely-moving rats using fiber photometry (Extended Data Fig. 1B), which allows for in vivo monitoring of neuromodulator signals with sub-second temporal resolution.

We tested the role of acetylcholine in learning across four distinct experiments using fiber-photometry recording or optogenetic manipulation. All experiments used the same odor-guided Go/No-Go trial structure and differed only in odor identity and reward contingency (Fig. 1A). In each trial, rats sampled an odor presented at a centrally located port and then decided whether to respond at a nearby fluid well based on the presented odor. Responses to rewarded cues resulted in a water reward, while responses to non-rewarded cues resulted in no reward and timeout penalties.

**Fig. 1.**
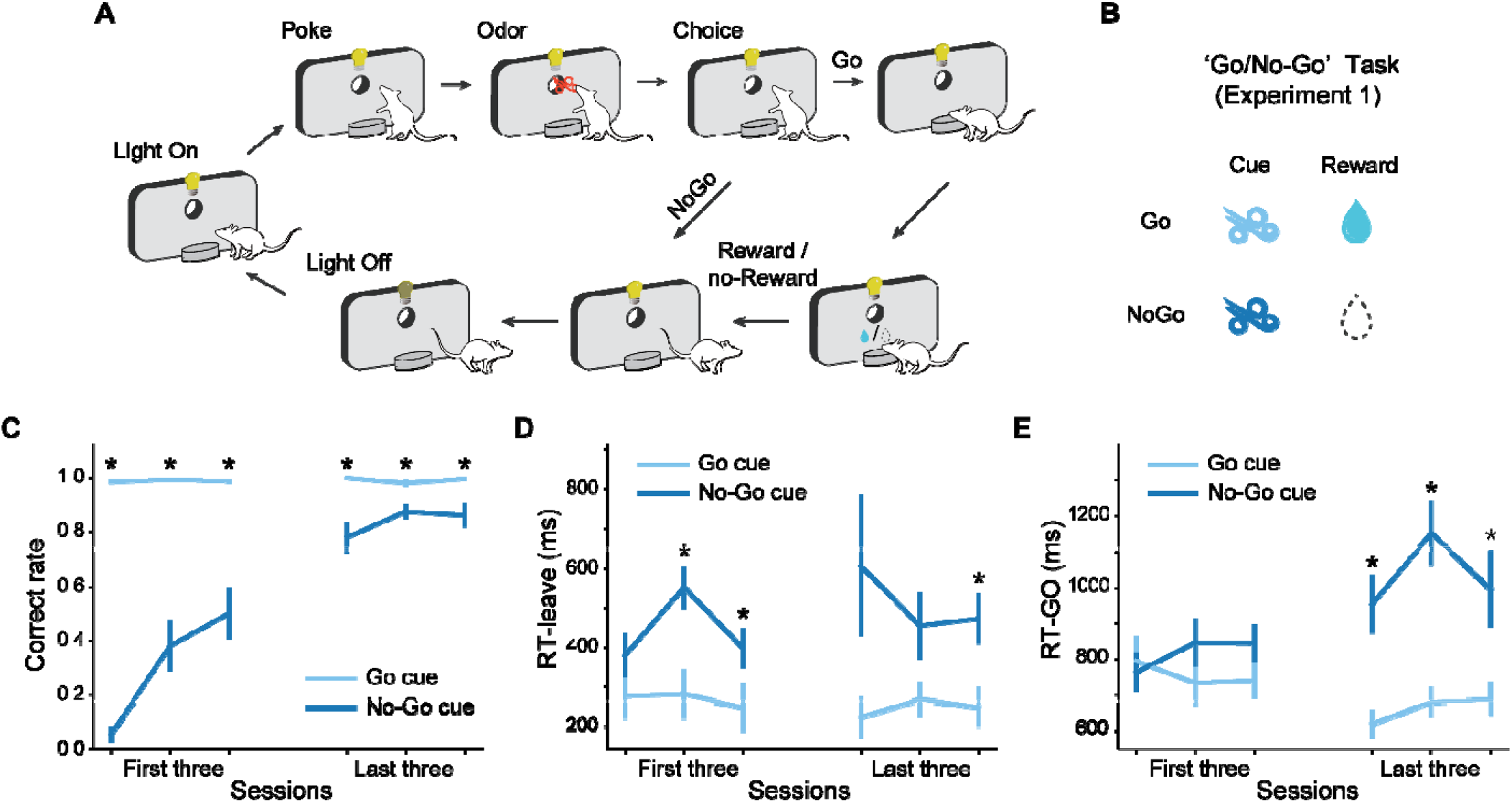
Go/No-Go task structure and behavioral performance. (**A**) Schematic diagram of the basic structure during a trial across the backbone Go/No-Go contingency used in the four tasks applied in this study. (**B**) Reward contingency for Experiment 1. Two cues are used in the Go/No-Go task. The Go cue indicates a water reward at the fluid well, while the No-Go cue indicates no reward. (**C**) Proportion of correct trials. The correct rates of No-Go trials (dark blue line) increased across training sessions and stayed high for the Go trials (light blue line). (**D**) Latency to leave the central port. Over learning, after the termination of odor delivery, rats left the central port quicker following the Go cue compared to the No-Go cue. (**E**) Latency to enter the fluid well. As learning progressed, rats became quicker at entering the fluid well after leaving the central port following the Go cue compared to the No-Go cue in the few trials where rats responded to the No-Go cue. The error bars in (C-E) indicate S.E.M. across rats. * indicates p < 0.05 in a two-tailed paired t-test.

### Acetylcholine and dopamine show distinct dynamics during discrimination learning

In Experiment 1, we explored the role of acetylcholine during associative learning in a Go/No-Go task with two distinct odors: a Go cue indicating a reward would be delivered and a No-Go cue indicating no reward (Fig. 1B). Initially, rats responded at the fluid well after presentation of each cue. Over time, they learned to withhold responses after the No-Go cue to avoid timeout penalties and maximize reward acquisition over time (Fig. 1C). Correct responses were defined as responding at the fluid well after the Go cue and not entering the fluid well after the No-Go cue. Rats were considered proficient when they reached 18 correct responses in a 20-trial moving block, which they accomplished on average within 285.38 ± 43.19 trials in 3.08 ± 0.55 sessions. In addition to increasing accuracy over the course of learning, the rats progressively decreased their response reaction time following the Go cue and increased it after the No-Go cue, both for leaving the odor port and entering the fluid well (Fig. 1D and E). These findings show that the rats successfully learned to discriminate between the two odor cues and associated each with the corresponding reward outcome.

Before turning to how acetylcholine and dopamine dynamics evolved during learning, we first characterized their responses in sessions when rats had already reached the behavioral criterion. Trials were grouped based on the presented odors. Consistent with dopamine’s role in representing RPEs, dopamine signals increased before odor onset and continued to rise after the representation of the Go cue, while they decreased following the No-Go cue (left panel, Fig. 2A). In contrast, the acetylcholine signals did not deviate from baseline until rats entered the odor port, as we previously reported ^14^, and were then reduced in response to both Go cue and No-Go cue (right panel, Fig. 2A). We quantified the cue-evoked response by averaging acetylcholine or dopamine signals between odor delivery and odor port exit, including the period after odor offset when the rat remained in the port, across all rats and sessions. As expected, dopamine responses were higher for the Go cue than the No-Go cue (left panel, Fig. 2B, *p* =2.2e-278 in a two-tailed t-test). In contrast, acetylcholine responses were significantly suppressed by both cues, with more pronounced dips for the Go cue (right panel, Fig. 2B, *p* =4.0e-30).

**Fig. 2.**
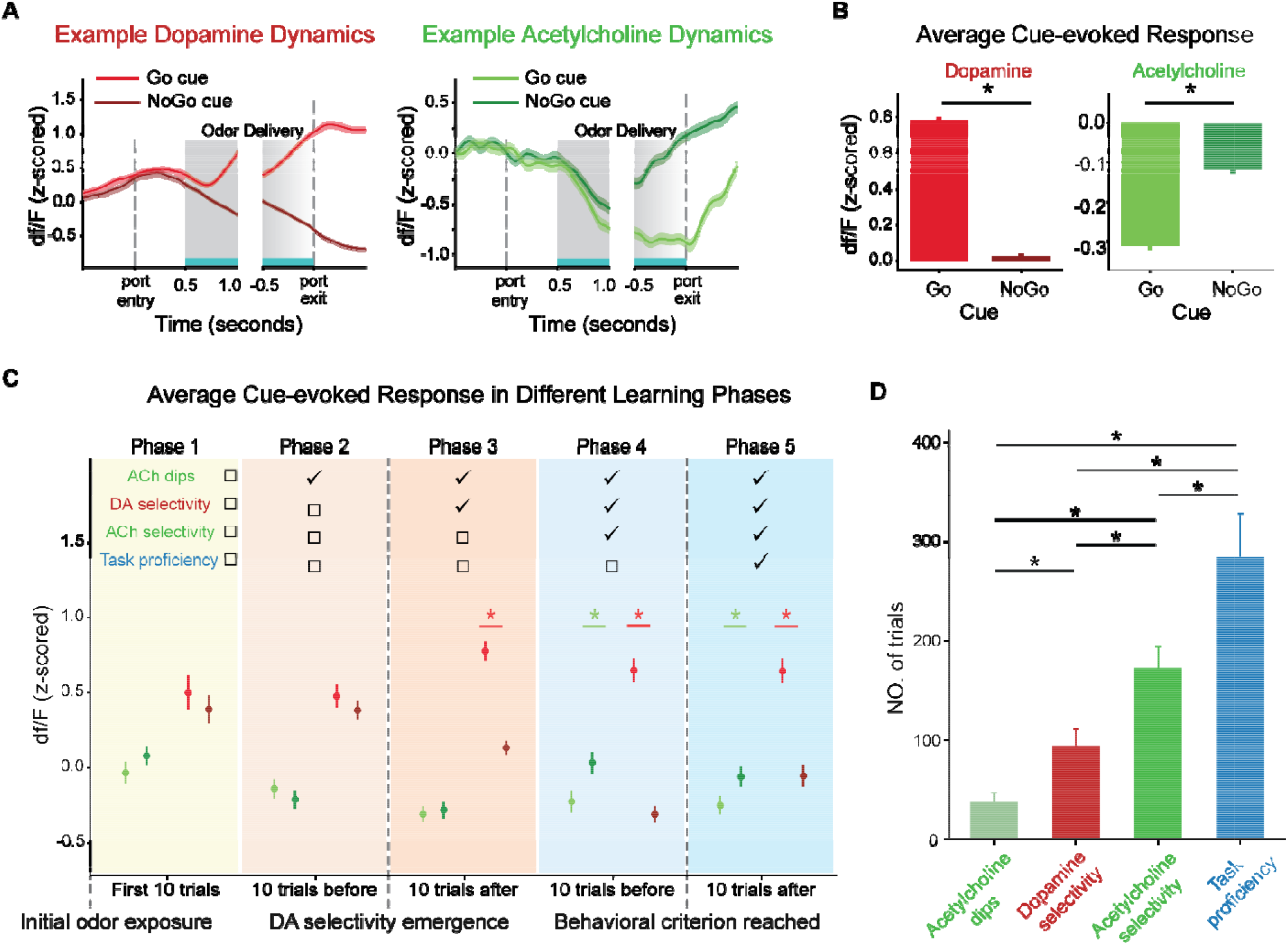
Dynamics of dopamine and acetylcholine during the Go/No-Go task (Experiment 1). (**A**) Example dopamine (left panel) and acetylcholine (right panel) dynamics within trials. The light and dark lines represent signals in the Go and No-Go trials, respectively. Dopamine responses increase over time in response to the Go cue and decrease during the No-Go cue trials, while acetylcholine responses drop during the odor delivery for both cue types. The gray shaded areas indicate the odor delivery period. The solid gray box marks the first 500 ms after odor onset, where odor delivery is present in all trials. The gradient gray box represents the last 500 ms before port exit, where odor presence varies. In trials where rats left the port quickly after odor offset, odor was still present, whereas in slower trials, it had already ceased. (**B**) Averaged cue-evoked dopamine (left) and acetylcholine (right) responses across all rats and sessions, computed during the period indicated by the blue bar on the *x*-axis in (A). Dopamine responses are significantly elevated during Go trials compared to the No-Go trials (=2.2e-278). Acetylcholine responses are significantly suppressed during both trial types and exhibit greater suppression during Go trials (=4.0e-30). (**C**) Averaged cue-evoked dopamine (left) and acetylcholine (right) signals across three learning milestones: learning initiation, the emergence of dopamine selectivity, and clear behavioral discrimination between cues. The error bars in (B and C) indicate S.E.M. across trials. Each * indicates p < 0.05 in a two-tailed t-test. (**D**) The number of trials required for the emergence of acetylcholine dips, dopamine selectivity, acetylcholine selectivity and reaching the behavioral criterion. The emergence of acetylcholine dips occurs first, followed by dopamine selectivity, then acetylcholine selectivity, and finally, the achievement of the behavioral criterion. The error bars in (D) indicate S.E.M. across rats. Each * indicates p < 0.05 in a Bayesian bootstrap test.

To investigate how these contrasting patterns of dopamine and acetylcholine release were established over learning, we divided the sessions leading up to this point into 5 phases defined by three key behavioral and neural milestones. The first milestone occurred when rats were initially exposed to the odor cues and had no prior knowledge of potential associations between the odor cues and reward. The second milestone was when dopamine responses to Go and No-Go cues first became significantly different across 10 successive trials (paired two-tailed t-tests, *p* <0.01). This served as a neural correlate of the learning that odors have different associations, even though behavioral differences between Go and No-Go cues might not yet be apparent at this stage. The third milestone was when rats showed a clear behavioral distinction between Go and No-Go cues, quantified as when the correct rate first reached 90%.

Phase 1 comprised the first 10 trials immediately following the first milestone, which was initial odor exposure. In this phase, dopamine responses were similar regardless of cue identity, consistent with the two-tailed t-test, *p* =0.49). Acetylcholine responses were also statistically indistinguishable from zero for idea that the specific cue-outcome associations had not yet been established (left panel, Fig. 2C. paired both Go and No-Go cues as expected (right panel, Fig. 2C. one sample t-test, *p* =0.66 and 0.63 for Go and No-Go trials, respectively). Phases 2 and 3 included the 10 trials immediately preceding and significant acetylcholine dips to both cues could be observed (*p* =9.2e-6 and 8.1e-4 for Go and No-Go following the second milestone, which was the emergence of dopamine selectivity. In phase 2, trials, respectively), but with no significant difference between them (*p* =0.27). In phase 3, although significant cue selectivity (*p* =0.37), which means that differentiation of dopamine cue responses dopamine responses discriminated between Go and No-Go cues, acetylcholine dips still did not show preceded differentiation of acetylcholine cue responses. Phases 4 and 5 included the 10 trials both phases, we found that acetylcholine dips showed significant cue selectivity (*p* =7.5e-6 and 3.9e-4, immediately preceding and following the third milestone, when rats reached behavioral proficiency. In respectively). Representative acetylcholine and dopamine PSTHs in different stages are shown in Extended Data Fig. 2.

To further quantify the temporal progression of cue-evoked acetylcholine and dopamine responses, we analyzed when acetylcholine responses deviated from baseline and when acetylcholine and dopamine responses differentiated between Go and No-Go cues over successive 10-trial bins. Our results revealed a sequence of emergence: cue-evoked acetylcholine dips, defined as a significant drop below zero, appeared first at approximately 38.15 ± 9.00 trials (mean +/-S.E.M. across rats), followed by dopamine selectivity, defined as differential responses to Go versus No-Go cues, at 94.46 ± 16.59 trials (*p* =2.3e-4, later, at 172.77 ± 21.79 trials (*p* < 1e-5 and =6.9e-4 compared to acetylcholine dip and dopamine Bayesian bootstrap test, see Methods; Fig. 2D). Acetylcholine selectivity to cues emerged significantly selectivity, respectively), and the behavioral criterion was reached last, at 285.38 ± 43.19 trials (*p*<1e-5, =5e-5 and 0.011 compared to acetylcholine dip, dopamine selectivity and acetylcholine selectivity, respectively).

This sequential evolution highlights a structured progression during learning: non-selective dips in acetylcholine emerge first, followed by dopamine selectivity that marks the differentiation of cue-outcome associations. Selective differences in acetylcholine dips appear after that, and ultimately, stable behavioral learning is achieved. This temporal hierarchy provides insight into how cholinergic and dopaminergic systems evolved dynamically during learning.

### Acetylcholine correlates with dopamine response updates and changes in behavior

The suppression of acetylcholine by both Go and No-Go cues, along with the delayed emergence of acetylcholine selectivity compared to dopamine selectivity, supports the idea that acetylcholine is not directly involved in value or RPE representation. Rather, these dynamics suggest that acetylcholine could contribute to acquired or associative salience ^12,13^, a major contributor to the rate at which associations are learned about specific cues

When rats were initially exposed to the odor cues and associative salience was minimal (Phase 1), cue-evoked acetylcholine responses were statistically indistinguishable from zero. According to this interpretation, the early acetylcholine dip preceding dopamine selectivity would reflect initial attentional engagement with the cues, a prerequisite for learning (Phases 2 and 3). The later emergence of acetylcholine selectivity would indicate a refined assignment of salience to specific cues based on their learned value (Phases 4 and 5), suggesting a complementary role of acetylcholine to the function of dopamine in learning cue-specific associations.

If cue-evoked acetylcholine dynamics indeed signal associative salience, then the magnitude of acetylcholine dips should correlate with changes in both dopamine responses and behavior and be a reliable predictor of learning rate. To examine this, we asked whether the magnitude of acetylcholine dips was associated with the rate at which outcome-evoked prediction errors updated cue value and behavioral choices.

Given the role of dopamine signals in representing RPE, we used reward-evoked dopamine responses to estimate the reward-evoked prediction error, and changes in cue-evoked dopamine responses across trials to estimate cue value updates. To test whether the acetylcholine dips correlate with the rate of cue value updating, we performed vector autoregression analyses in which four sets of predictors, i.e., cue-evoked acetylcholine dips, reward-evoked dopamine response, their interaction term and the changes in cue-evoked dopamine from the previous *N*_lag_ trials, were used to predict estimated cue value updates. The lag length, *N*_lag_ = 1, was determined in advance using Bayesian information criteria (Extended Data Fig. 3A). Using Granger causality tests, we observed a significant effect of reward-evoked dopamine on cue value updates compared to shuffled controls (two-tailed t-test, *p*=4.5e-8, Fig. 3A), consistent with the role of dopamine in representing RPE. Importantly, a strong and significant effect of the interaction between reward-evoked dopamine and cue-evoked acetylcholine responses on cue value updates was also detected (*p*=1.3e-6, Fig. 3A), confirming that acetylcholine dips correlated with the updating of cue value.

**Fig. 3.**
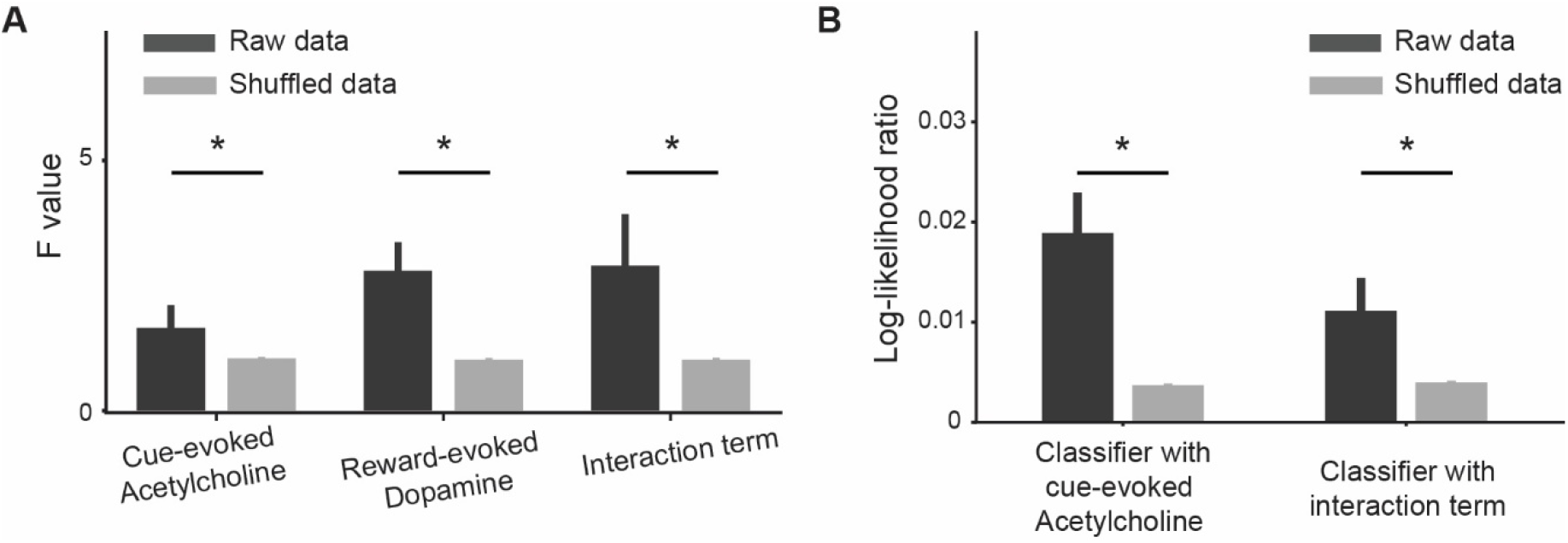
Acetylcholine release correlates with behavior and dopamine updates. (**A**) F-values from the Granger causality test for the actual data (dark grey bars) versus shuffle data (light grey bars). Significant effects were found for reward-evoked dopamine responses (=4.5e-8), its interaction with cue-evoked acetylcholine responses (*p* =1.3e-6) on the subsequent cue-evoked dopamine responses, and the cue-evoked acetylcholine responses (=0.039). (**B**) The log-likelihood ratio of logistic classifiers that included either acetylcholine (acetylcholine model) or its interaction with dopamine against a baseline classifier using only reward-evoked dopamine. Bars show the log-likelihood ratio values for actual data (dark grey) and for shuffled permutations (light grey). In both classifiers, the actual-data log-likelihood ratio is significantly higher than the shuffled distribution (p = 7.9e-18 for acetylcholine; p = 4.0e-5 for interaction). The error bars indicate S.E.M. across rats. Each * indicates p < 0.05 in a two-tailed t-test.

We then measured whether acetylcholine dips were associated with behavioral changes in the No-Go cue, excluding trials with the Go cue, as rats consistently responded to it throughout learning, leading to little variability. We first trained a logistic classifier to predict the behavior only based on the reward-evoked prediction error from the previous trial, also estimated by the reward-evoked dopamine response. Next, we constructed two extended classifiers, one including the cue-evoked acetylcholine dips and another including the interaction term as additional predictors. The contribution of each predictor was evaluated by the log-likelihood ratios (logLR) between extended classifiers and the basic one. To evaluate the significance, we shuffled the cue-evoked acetylcholine dips and interaction term 100 times and recalculated the logLR for each shuffle. In both cases, classifiers with actual data showed significantly significantly higher logLR compared to the shuffled controls, indicating that the acetylcholine signal (*p*=7.9e-18, two-tailed t-test, Fig. 3B) and its interaction with prediction error (*p*=4.0e-5) contributed to behavioral predictions. These findings demonstrate that the cue-evoked acetylcholine dips correlate with the rate of cue value updating and behavioral adaptation, supporting the hypothesis that acetylcholine responses encode associative salience and modulate learning rate.

### Optogenetic manipulation of accumbal cholinergic interneurons affects the rate of associative learning

The correlational analyses described above showed that cue-evoked acetylcholine dips were associated with both dopamine response updates and behavioral changes. We next tested whether cue-evoked acetylcholine release contributes causally to the learning process, consistent with a role in signaling associative salience. To test this, we manipulated acetylcholine release in the NAcc by optogenetically increasing or decreasing the activity of accumbal cholinergic interneurons, the primary source of acetylcholine in this region, while assessing behavioral learning in rats.

Twenty ChAT-Cre rats were assigned to one of three groups: activation, inhibition, or control. Rats in the activation group (n = 5) received bilateral NAcc injections of the Cre-dependent viral vector pAAV-Ef1a-DIO-hChR2-EYFP, rats in the inhibition group (n = 5) received pAAV-Ef1a-DIO-eNpHR-EYFP, and control rats (n = 10) received pAAV-Ef1a-DIO-EYFP. Bilateral optic fibers were implanted above the injection sites for light delivery. Fiber placement and viral expression were verified histologically and by immunostaining (Fig. 4A, Extended Data Fig. 1A), with EYFP expression evaluated in relation to VAChT immunoreactivity. Behavioral learning was assessed in four sequential Go/No-Go problems, each involving a unique odor pair (Experiment 2, Fig. 4B). Training for each problem was stopped 20 trials after rats achieved the behavioral criterion. Rats were then progressed to the next problem in the subsequent session until they had learned all four problems. During cue presentation, activation rats received 465 nm blue light, inhibition rats received 590 nm yellow light, and control rats were evenly matched to these light-delivery protocols. In all groups, light delivery began 100 ms before odor onset and ended at odor offset, ensuring that optogenetic manipulation fully covered the cue presentation period.

**Fig. 4.**
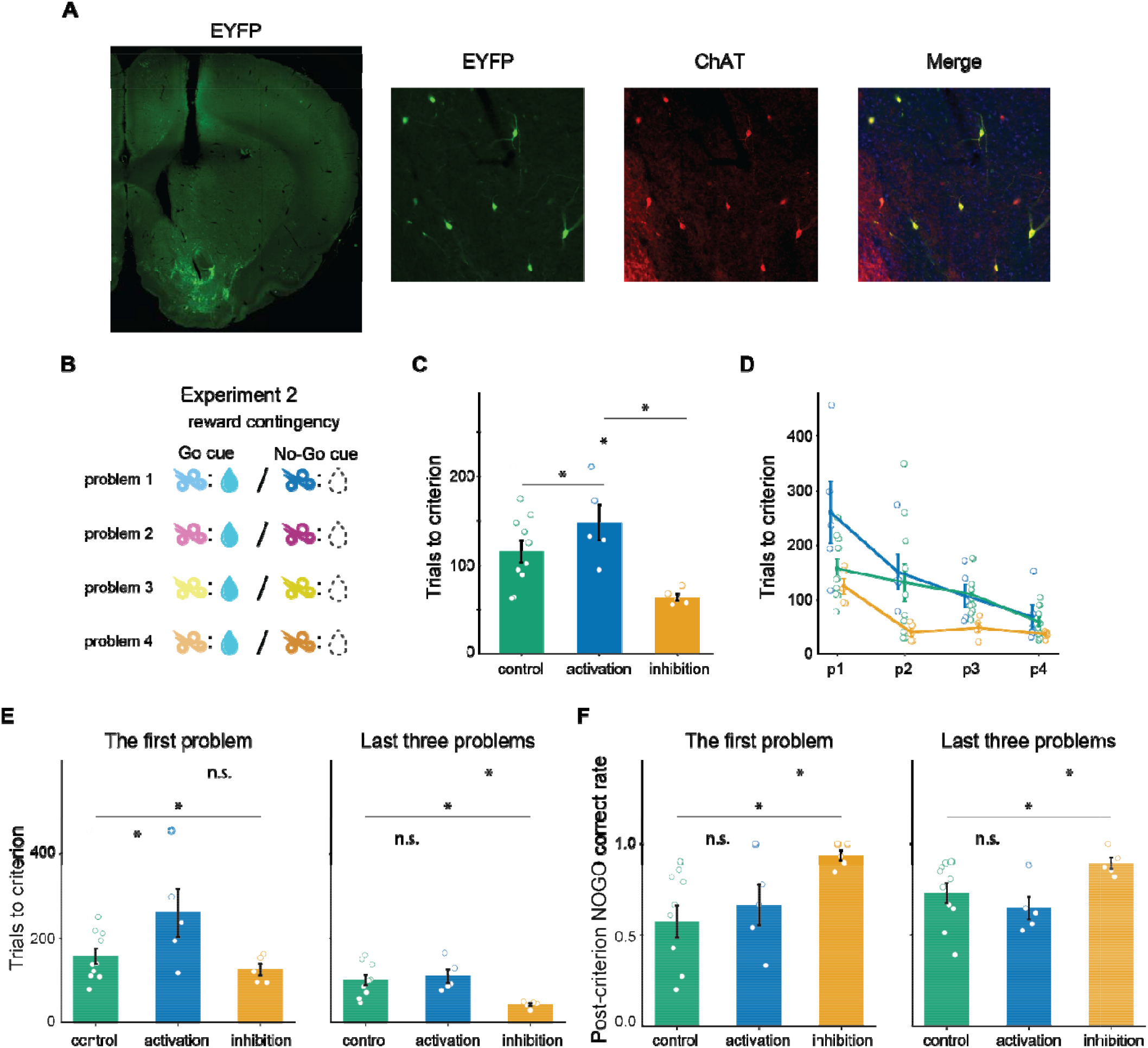
Optogenetically manipulating cholinergic neurons bidirectionally modulates learning speed in associative learning. (**A**). Virus expression and colocalization with VAChT. A represent section shows viral expression in the NAcc core following injection of an AAV encoding a Cre-dependent EYFP-tagged optogenetic construct. EYFP expression was visualized by anti-GFP immunohistochemistry (green), and cholinergic neurons were labeled with an anti-VAChT antibody (red), with nuclei counterstained with DAPI (blue). (**B**) Successive Go/No-Go problems, each involving a unique pair of odor cues. (**C**) Mean number of trials required to reach the behavioral criterion, averaged across all four problems for each rat. Each data point represents a rat. (**D**) Trials required to reach the behavioral criterion for each group and problem. **(E)** Trials required to reach criterion in the first problem (left panel) and in later problems (right panel). Optogenetic activation increased trials to criterion in the first problem, whereas optogenetic inhibition reduced trials to criterion in later problems. **(F)** No-Go correct rate during post-criterion trials in the first problem (left panel) and in later problems (right panel). Inhibiting accumbal cholinergic interneurons increased post-criterion No-Go accuracy in both first and later problems, indicating inhibition facilitated learning and improved post-criterion performance. In all panels, dots indicate individual rat values and error bars indicate mean ± S.E.M. Pairwise group comparisons were evaluated using Bayesian bootstrap tests, and * indicates p < 0.05.

As above, we quantified learning performance as the number of trials required for each rat to reach the behavioral criterion. For each rat, we first averaged the number of trials to criterion across all four problems and compared these values across optogenetic groups (Fig. 4C). Rats in the activation group required an average of 147.6 trials to reach criterion, whereas control rats required 115.2 trials, 32.4 fewer than activation rats. We assessed group differences using Bayesian bootstrap analysis, which showed that rats in the activation group required significantly more trials to reach criterion than control rats (p = 0.044). In contrast, rats in the inhibition group required 63.6 trials, which was 51.6 fewer than controls (p < 1.0e-4) and 83.9 fewer than activation rats (p < 1.0e-4). These results suggest that reducing accumbal cholinergic interneuron activity, thus enhancing acetylcholine dips, during cue presentation facilitated associative learning, whereas activating these interneurons, thus preventing acetylcholine dips, slowed associative learning.

Interestingly, the effects of cholinergic manipulations also covaried across successive learning problems. Inspection of the trials to criterion across problems (Fig. 4D) showed that the disruptive effect of activation was most evident on the initial problems, while the facilitative effect of inhibition was most evident in the later problems. To quantify these problem-dependent effects, we repeated the Bayesian bootstrap analysis for the first problem and the later problems separately. Results showed that the activation of accumbal cholinergic interneurons increased trials to criterion in the first problem compared to controls (p value = 8.8e-3; left panel, Fig. 4E), which, however, was not observed in the later problems (p value = 0.31; right panel, Fig. 4E). In contrast, optogenetic inhibition reduced trials to criterion in the later problems compared to controls (p value < 1.0 e-4). In the first problem, inhibition showed a trend toward reducing trials to criterion, directionally consistent with the observation in later problems, although this effect did not reach significance (p value = 0.053). As an additional measure of learning performance, we quantified the post-criterion No-Go correct rate. Higher rates indicated that more accurate and stable performance had been reached. Consistent with the above finding, rats in the inhibition group showed a higher post-criterion No-Go correct rate than controls in the first problem (p value < 1.0e-4; left panel, Fig. 4F). This enhancement in post-criterion No-Go performance was also observed in the later problems (p value < 1.0 e-4; right panel, Fig. 4F). These temporal effects suggest that the cholinergic dips may be particularly critical when rats first associate any odor cues with motivational events within a particular task context. Under these conditions, the dips may play a particularly critical role in adjusting the salience and learning rate to the appropriate cues.

Overall, these findings further support the hypothesis that cue-evoked acetylcholine dips determine the learning rate, which is fundamentally dependent on associative salience. Thus, bidirectional manipulation of accumbal cholinergic interneurons provided causal evidence that cue-evoked acetylcholine modulates learning rate during associative learning.

### Acetylcholine dynamics reflect cue-related salience changes after reward contingency reversal

The above experiments and analyses demonstrate that cue-evoked acetylcholine dips in a stationary environment correlate with and modulated learning rate in a manner that mimics an associative salience signal. In Experiment 3, we aimed to extend our findings by testing if acetylcholine dips track salience signals in a dynamic environment where reward contingencies change over time. Previous studies have shown that the salience of cues does not automatically update following changes in reward value^29,30^, and the salience of a cue can persist even after decreases in the value of its predicted outcome ^31^ as occurs in reversal tasks. According to this, we hypothesized that after reversal in a Go/No-Go task, the acetylcholine dips to the previously rewarding cue would persist, while the dips to the previously non-rewarded cue would increase in magnitude due to its new association with reward.

To test these predictions, we performed Experiment 3, which consisted of a series of three reversal problems (Fig. 5A), each involving two distinct stages. In stage 1, conditioned stimulus 1 (CS1) was rewarded, while conditioned stimulus 2 (CS2) was non-rewarded. Training continued for an additional 20 trials after rats first achieved the behavioral criterion. Beginning in the next session, the original reward contingencies remained until rats met the behavioral criterion again, at which point we initiated stage 2 and reversed the reward contingencies associated with CS1 and CS2. Training with these reversed contingencies continued across multiple sessions until rats met the behavioral criterion again. After successfully completing the first reversal task, we repeated this two-stage process with a second and third set of odor pairs.

**Fig. 5.**
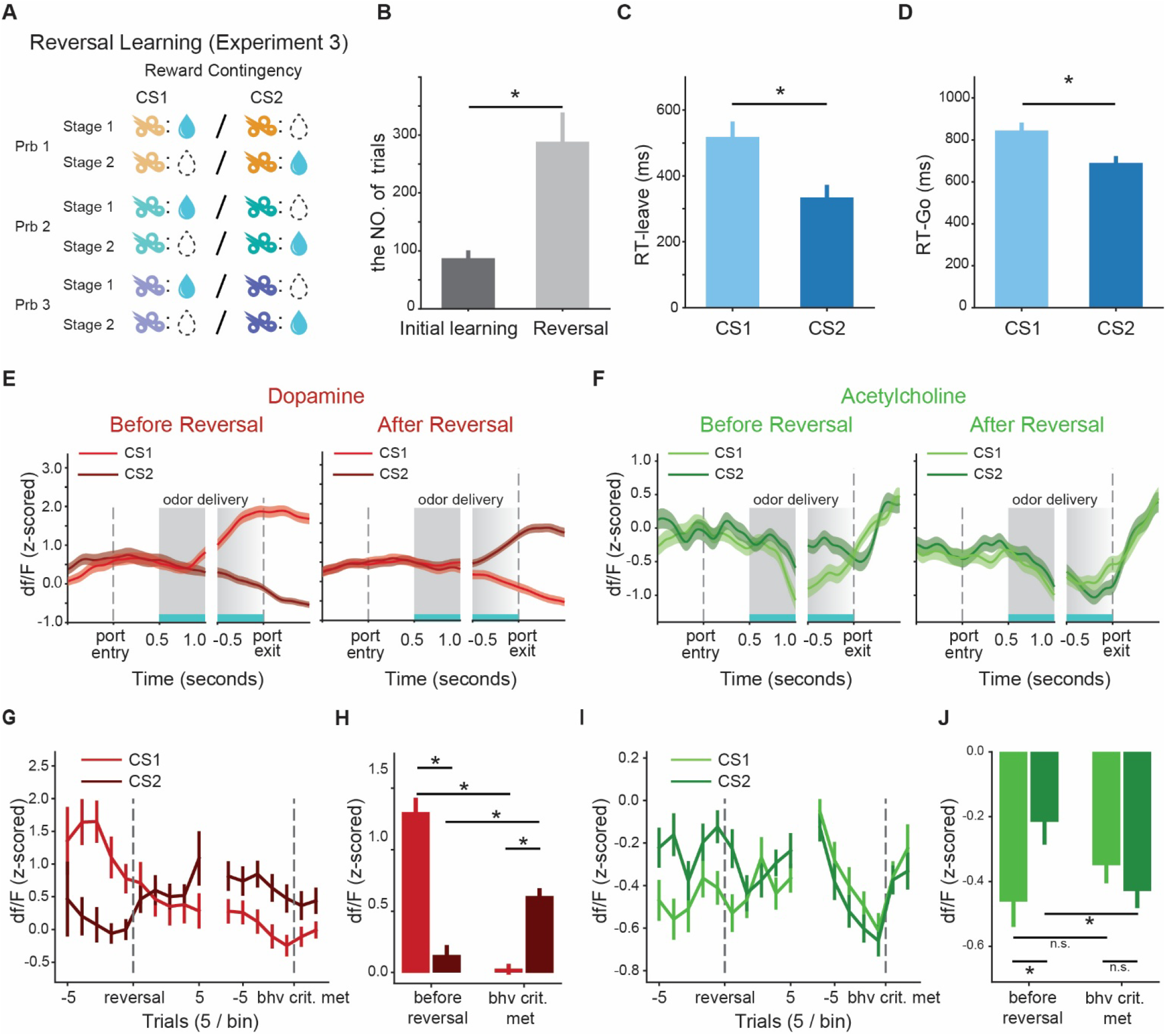
Acetylcholine responses to the previously rewarded cue persisted after reversal (Experiment 3). (**A**) Schematic of reward contingencies in Experiment 3. The reversal learning task is composed of three problems, each with a different odor pair. The reward contingencies are reversed during the learning. (**B**) Number of trials required to reach the behavioral criterion during initial training (Stage 1) and after the reversal (Stage 2). (**C**) Latency to leave the odor port following CS1 or CS2 presentation after the reversal. (**D**) Latency to enter the fluid well after CS1 or CS2 after the reversal. Rats exhibited shorter latencies for CS2 compared to CS1 after reversal. (**E**) Example dopamine dynamics before (left panel) and after reversal (right panel) within trials. (**F**) Example acetylcholine dynamics before (left panel) and after reversal (right panel) within trials. (**E-F**) follow the same format as Fig. 2A, respectively. (**G**) Dynamics of the dopamine responses around the reversal point and upon reaching the reversal learning criterion are shown for CS1 and CS2. (**H**) Average dopamine responses to cues before the reversal and upon meeting the post-reversal behavioral criterion. (**I** and **J**) Acetylcholine signal dynamics and average responses. These panels follow the same format as panels (E and F), respectively. * in (**B**-**D, H**, and **J**) indicates <0.05 in a two-tailed t-test.

Rats reached the post-reversal behavioral criterion with training, but required more trials than the initial reward contingency learning (Fig. 5B; 272.6 ± 44.8 vs. 95.3 ± 15.3 trials, respectively; paired two-tailed t-test, *p*=0.0011). After reversals, rats exhibited shorter response latencies in both leaving the odor port (Fig. 5C, *p*=5.8e-4) and entering the fluid well (Fig. 5D, *p*=5.1e-5) following CS2 compared to CS1. These behavioral changes suggested that after reversal, rats associated both a higher value and greater motivation with CS2.

To gain an overall view of how dopamine and acetylcholine signals changed dynamically during reversal learning, we plotted the PSTHs of dopamine and acetylcholine signals from a representative rat, averaging across trials after the behavioral criterion was achieved both before and after reversal. Before reversal, dopamine responses were higher for CS1 compared to CS2, and this pattern reversed after reaching the post-reversal criterion (Fig. 5E). In contrast, acetylcholine showed a different pattern, initially exhibiting larger dips to CS1 before reversal. However, after the behavioral criterion was met, acetylcholine dips to CS1 persisted and the acetylcholine dips to CS2 were enhanced (Fig. 5F).

To further quantify these dynamic changes across all rats, we averaged these signals around the reversal point and upon reaching the reversal learning criterion. The dynamics of the dopamine response remained consistent with RPEs throughout the task. Before reversals, the cue-evoked dopamine response to CS1 was significantly larger than that to CS2 (*p*=6.2e-13, left side of Fig. 5 G and H). After meeting the post-reversal criterion, dopamine responses to CS2 increased (*p*=1.1e-5) and the responses to CS1 decreased compared to pre-reversal levels (*p*=9.9e-23, Fig. 5 G and H). As a result, the dopamine responses to CS2 became significantly higher than CS1 (right side of Fig. 5 G and H, *p*=1.1e-12).

We applied the same analysis to acetylcholine responses. Consistent with Experiment 1, before reversals, both odors suppressed the acetylcholine signal, with CS1 inducing a more pronounced dip compared to CS2 (left side of Fig. 5I, *p*=0.02). After the reversal, the magnitude of the acetylcholine dip evoked by CS2 increased while the dip evoked by CS1 persisted. In contrast to the dopamine responses, when rats reached the new criterion, acetylcholine dips to CS2 increased (*p*=0.01), while the acetylcholine dips to CS1 showed a slightly non-significant decrease (*p*=0.23, Fig. 5 I and J). Consequently, acetylcholine dips to CS1 and CS2 were statistically indistinguishable (right side of Fig. 5 I and J, *p*=0.30). These results are consistent with our prediction that the acetylcholine dips to the previously rewarding cue (CS1) remained for an extended period, while dips to the newly rewarding cue (CS2) were enhanced. This supports the view that acetylcholine dips track a salience signal in dynamic environments and update more slowly than the value. Furthermore, the lack of differential acetylcholine responses, despite rats showing higher motivation toward CS2 (Fig. 5 C and D), argues against the interpretation that acetylcholine primarily reflects motivation or value.

### Acetylcholine dynamics and selectivity in the Go/No-Go task are captured by a hybrid attentional associative learning model

To quantitatively assess whether acetylcholine dips are consistent with salience signaling, we first analyzed acetylcholine dynamics and then compared them to the normative salience terms defined by different attentional associative learning models. We initially examined the cue-evoked acetylcholine responses for each odor cue condition over the first 150 trials in the Go/No-Go task (Experiment 1). A biphasic trend was observed for the No-Go cue: the magnitude of acetylcholine dips initially increased, followed by a subsequent decrease (Fig. 6A). This decreasing trend held when we averaged the the subsequent sessions compared to the first session (*p*=2.0e-5, 0.038, and 6.2e-4 compared to the responses per session, with the No-Go cue-evoked acetylcholine dips significantly less pronounced in 2^nd^/3^rd^/4^th^ sessions, respectively, Fig. 6B). In contrast, acetylcholine dips to the Go cue gradually increased in magnitude over learning and showed little evidence of subsequent decline (Fig. 6 A and B), resulting in consistently stronger dips compared to the No-Go cue. We also analyzed the dopamine responses in the same way, and the results (Extended Data Fig. 4) were consistent with RPE dynamics.

**Fig. 6.**
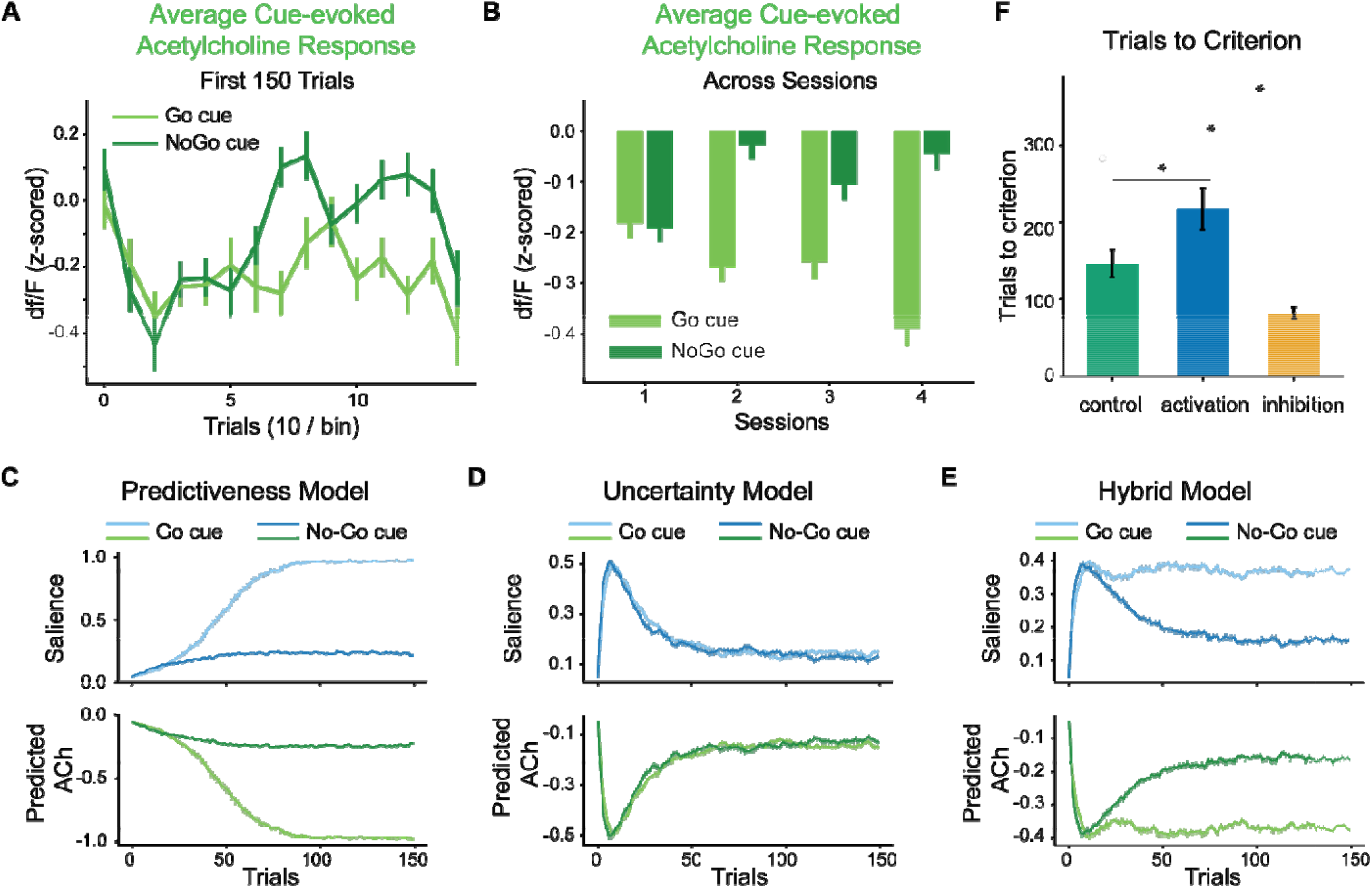
A hybrid attentional associative learning model captures biphasic dynamics and cue selectivity of the acetylcholine responses. (**A**) Average cue-evoked acetylcholine responses for Go (light green) and No-Go (dark green) cues in the first 150 trials. (**B**) Averaged cue-evoked acetylcholine responses for Go (light green) and No-Go (dark green) cues across training sessions. The error bars indicate S.E.M. across trials. (**C-E**) The upper panels show the dynamics of salience predicted by the predictiveness model (**C**), based on the Mackintosh model; the uncertainty model, based on the Pearce-Hall model (**D**); and the hybrid model (**E**), which integrates salience mechanisms from both models. The lower panels show the acetylcholine signal predicted by the models. The light lines indicate the Go cue, while the dark lines indicate the No-Go cue. The error bars indicate S.E.M. across 100 simulations with different random seeds. (**F**) Simulated optogenetic modulation alters the number of trials required to reach the learning criterion. Bars show the mean number of trials required for the hybrid model to reach 90% correct performance under simulated activation, control, and inhibition conditions. The error bars indicate S.E.M. across 20 simulations. * indicates <0.05 in a two-tailed t-test.

Subsequently, we reconstructed the task using computational models that reflect the attentional associative learning framework^32-35^, where salience modulates the learning rate and, together with RPEs, adjusts stimulus-outcome associations. The most influential models in this framework are the Mackintosh ^32,36^ and Pearce-Hall models ^35,37^. Critically, each model proposes a different definition of salience, which leads to different predictions. Mackintosh’s model assigns high salience to cues that are more predictive of outcomes than others, referred to as predictiveness-driven salience (PDS). The PDS is further modulated by the absolute outcome strength, reflecting the fact that cues predicting higher reward value or stronger punishment retain greater salience. This model explains various phenomena, such as the observation that previously blocked cues are slower to support novel associations^38,39^. In contrast, the Pearce-Hall model defines salience as the mean absolute value of the RPE over time, reflecting the uncertainty in predicting outcomes. This uncertainty-driven salience (UDS) reflects the fact that animals allocate more attention to cues with uncertain outcomes. This model accounts for phenomena like latent inhibition^40,41^, where repeated exposure to an isolated cue impairs subsequent learning about associations to that cue. Hybrid models ^23,34^, which integrate salience components from both Mackintosh and Pearce-Hall models, have been proposed to conciliate these two views, and can determine a cue’s ultimate total salience, explaining a broader range of experimental findings. We built models (see Methods for details of all models) grounded in these theoretical accounts and performed 100 simulations with different random seeds for each to ensure robustness. As acetylcholine signals dipped in response to cues, we inverted the sign of the predicted salience and compared it with the acetylcholine response.

To explain the acetylcholine dynamics, we first applied the predictiveness model, grounded in the Mackintosh model, and the uncertainty model, grounded in the Pearce and Hall model. Each accounted for part of the results, but neither reproduced both the biphasic trend and cue identity selectivity. In the predictiveness model, both cues gained salience because they were the most reliable predictors among all perceivable stimuli (Fig. 6C). The Go cue was assigned a higher salience boundary since the Go cue was associated with a more motivationally salient outcome, i.e., water reward, compared to the No-Go cue’s outcome, i.e., reward omission or timeout penalty. However, this model predicted a monotonic increase in salience and therefore a monotonic decrease in the acetylcholine responses (Fig. 6C), which conflicted with the observed biphasic trend (Fig. 6A). In the uncertainty model, the associative salience depended on the outcome uncertainty. As the associations between cues and outcomes were learned, the RPE and predicted salience initially increased and then decreased (Fig. 6D). Thus, this model predicted identical salience patterns for both cues, especially once the associations were established (Fig. 6D), and failed to capture the acetylcholine selectivity in cue identity during learning.

Finally, we built a hybrid model which integrates mechanisms from both the Mackintosh and Pearce-Hall models of salience ^23^. In this model, salience was defined as the weighted sum of the PDS from the Mackintosh model and the UDS from the Pearce-Hall model. The PDS component increased and was modulated by the learned value of cues, mirroring the predictiveness model. In parallel, the UDS component showed a biphasic pattern during learning, similar to the pure uncertainty model. As a result, the salience signal in the hybrid model captured both the biphasic dynamics and cue selectivity of acetylcholine dips we observed in our experiments (Fig. 6 A and E).

To rule out the possibility that the distinct patterns of salience described for the models above were due to the arbitrarily chosen parameters, we fitted models by minimizing the mean squared error in reproducing acetylcholine responses with rat behavioral data. This optimization was performed using differential evolution. Due to the simulation’s complexity, we fitted learning rates for the association strength, PDS and UDS, which we believe most significantly affect the prediction error. The results revealed a significantly smaller mean square error of reproducing the acetylcholine responses by the hybrid model compared to the predictiveness model (*p*=3.7e-27, two-tailed t-test) and the uncertainty model (*p*=5.5e-18). Thus, the hybrid model comprehensively explained the observed acetylcholine response patterns, indicating that acetylcholine responses encode the combined predictiveness-driven and uncertainty-driven salience as proposed in Mackintosh and Pearce-Hall models.

We next asked whether associative salience from the hybrid attentional model could also account for the behavioral effects of bidirectional optogenetic manipulation of cholinergic activity. We modeled the effects of optogenetic manipulation by shifting model-derived salience in response to corresponding changes in cue-evoked ACh signals. Specifically, optogenetic activation, which attenuated the cue-evoked ACh dip, was modeled by applying a constant negative bias to total salience. This manipulation delayed learning, increasing the number of trials required to reach the behavioral criterion from 145.9 ± 17.7 in the control condition to 217.0 ± 27.5 in the activation condition (*p*=6.6e-4, two-tailed *t*-test, Fig. 6F). In contrast, optogenetic inhibition, which enhanced the ACh dip, was modeled by adding a positive bias to total salience. This manipulation accelerated learning, reducing the number of trials to criterion to 82.1 ± 7.1 (*p*=7.2e-4, two-tailed *t*-test, Fig. 6F). Thus, without altering any other model components, bidirectional shifts in model-derived salience were sufficient to reproduce the corresponding effects of cholinergic activation and inhibition on learning rate. Incorporating experience-dependent salience baseline and bias approaching the fluid well, the extended hybrid model captures both the faster learning over problems and the problem-dependent effects of optogenetic manipulation (Method and Extended Data Fig. 5).

### The hybrid attentional associative learning model explains acetylcholine dynamics in reversal learning and predicts the cue-evoked acetylcholine responses with extended training

Our hybrid associative learning model also successfully captured the characteristics of the acetylcholine dips in the reversal learning task (Experiment 3). After reversals, all odor cues predicted the opposite reward outcome, leading to elevated RPEs and diminished predictiveness. Consequently, there was an increase in UDS and a decrease in PDS (blue arrows in Fig. 7, A and B). Once the values were updated, the stimuli regained PDS due to enhanced predictiveness and lost UDS owing to reduced uncertainty (purple arrows in Fig. 7, A and B). For CS1, the changes in PDS and UDS largely canceled each other, leading to little change in the overall salience (light lines in Fig. 7C). In contrast, CS2’s low pre-reversal PDS and its new association with water reward caused an increase in PDS, leading to larger total salience (dark lines in Fig. 7C).

**Fig. 7.**
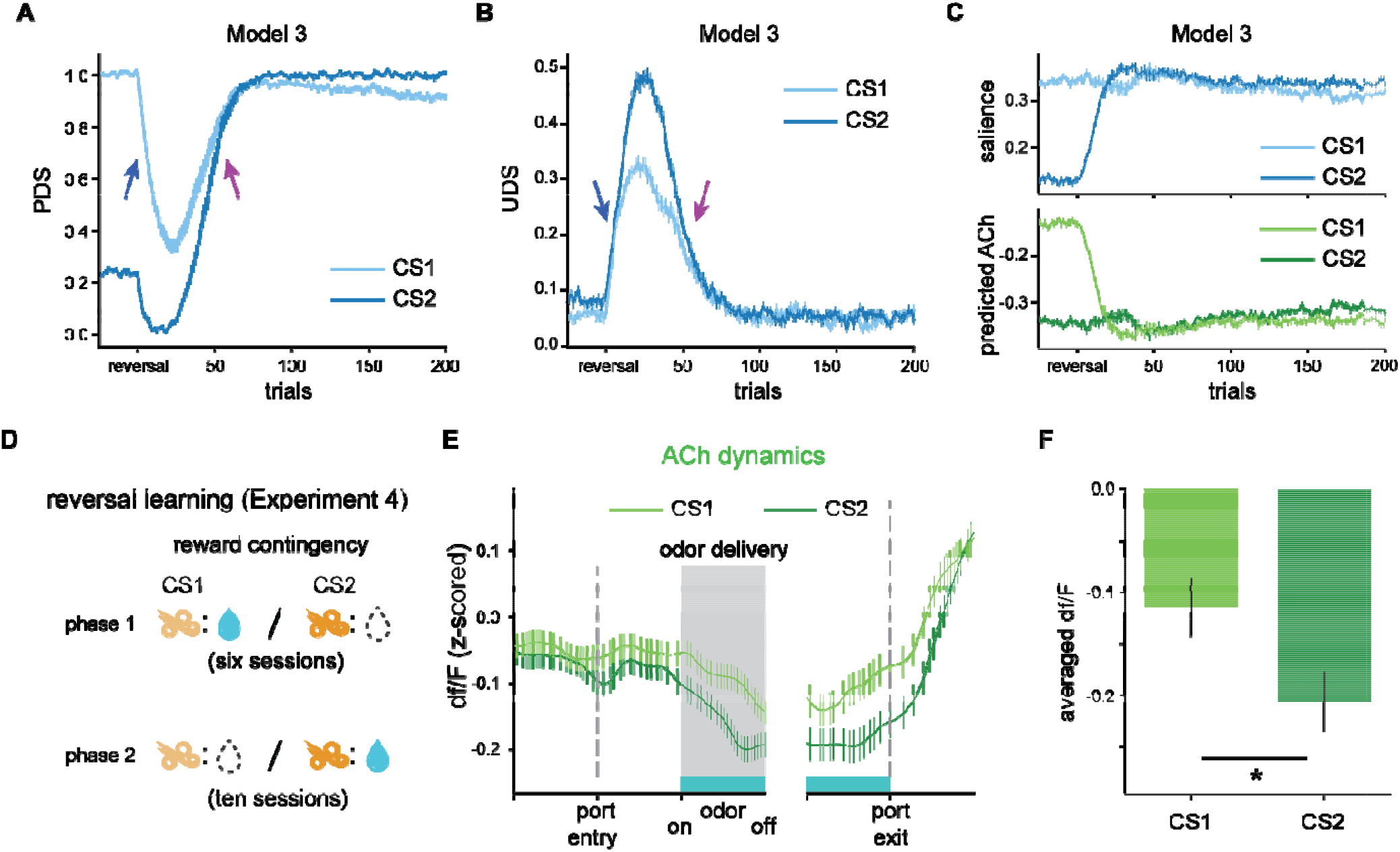
The hybrid associative learning model captured the dynamics of the acetylcholine signal in the reversal learning task. (**A-C**) Dynamics of predictiveness-driven salience (**A**), uncertainty-driven salience (**B**), total salience, and acetylcholine responses predicted (upper and lower penal in (**C**), respectively) by the hybrid associative learning model following the reversal of reward contingencies. The light and dark lines represent signals in the CS1 and CS2 trials, respectively. Initially, reversal leads to elevated RPEs, leading to increases in UDS and decreases in PDS (blue arrows). With learning, the stimuli regain PDS due to improved relative predictiveness and lose UDS as uncertainty declines (purple arrows). The PDS and UDS changes of CS1 largely cancel out, while the new reward-associated CS2 shows a net increase in total salience due to enhanced PDS following the value update. The error bars indicate S.E.M. across 100 simulations with different random seeds. (**D**) Schematic of reward contingencies in Experiment 4. (**E**) Empirical data showing the acetylcholine dynamics within trials after extended post-reversal learning. (**F**) Averaged cue-evoked acetylcholine responses across all rats in the last three sessions, showing the reversed cue selectivity in acetylcholine signaling. * indicates <0.05 in a two-tailed t-test.

The salience of CS1 remained high even when the post-reversal behavioral criterion was met due to the slow decline in the upper boundary of PDS, reflecting the persistent influence of reward history. However, the model does predict that, with extended training, this boundary would eventually decrease, causing low PDS and reduced total salience for CS1. Thus, we hypothesized that extended post-reversal training would invert the preference of acetylcholine dips in cues.

To test this prediction, we conducted Experiment 4 (Fig. 7D). In this experiment, five rats were trained in a Go/No-Go task for six sessions, and from the seventh session, we reversed the reward contingencies 20 trials after the rats achieved the behavioral criterion. Training with the reversed contingencies continued for ten sessions. Pooling the data from the final three sessions, we observed a reversed selectivity in acetylcholine dips. CS2 evoked significantly more pronounced acetylcholine dips than CS1 (*p*=0.025, Fig. 7 E and F). These results directly tested and confirmed a prediction of the hybrid model of associative salience, further supporting the proposal that acetylcholine dips encode the cue’s associative salience and demonstrating that their dynamics adapt to the changing reward contingencies over longer timescales.

## Discussion

Here, we explored the role of acetylcholine in the learning process by examining how acetylcholine signals evolve in the NAcc alongside dopamine signals and behavioral learning, across multiple experiments. We found that there were no acetylcholine responses to novel cues, but as cue-outcome associations were learned, characteristic cue-evoked dips in acetylcholine emerged. The magnitude of these dips changed over learning, and their selectivity for Go versus No-Go cues appeared later than dopamine selectivity but before rats reached behavioral proficiency. The magnitude of the acetylcholine dips also correlated with RPE-related dopamine transients and the behavioral learning rate, and temporally-specific manipulation of accumbal cholinergic interneurons during cue presentation altered learning rate, with activation slowing and inhibition facilitating associative learning. In addition, unlike dopamine responses, cue-evoked acetylcholine dips persisted even when the value of cues decreased, tracking the overall relevance of the cues, rather than their value. The lack of response to novel cues, the correlation with learning rate, the decoupling from value, and the unidirectional nature of these dynamic acetylcholine dips are most consistent with signaling of the associative salience acquired by the reward predictive cues. Accordingly, computational modeling showed that the cue-evoked acetylcholine responses were best explained by an acquired salience term that modulates learning rate, as defined by a hybrid attentional associative learning model that integrates salience driven by both predictiveness and uncertainty.

While many traditional reinforcement learning theories often assume a constant learning rate, optimal learning requires adjusting this rate based on environmental volatility^42,43^. Recent studies suggest that dopamine transients encode RPEs independently of learning rates^44^, raising the question of how the learning rate is determined. Previous studies have suggested that the cholinergic system is well-suited for this function. Extensive interactions between dopaminergic and cholinergic systems have been observed in multiple brain regions^16,22^, and acetylcholine dips in the NAcc have been proposed to create a permissive window for phasic dopamine increases to facilitate synaptic plasticity^5,45^, thus supporting the hypothesis that the acetylcholine dips could determine learning rates. Here, we directly demonstrated that acetylcholine in NAcc follows the salience changes predicted by a hybrid attentional associative learning model and affects the rate of dopamine and behavior updates, which is aligned with the role of salience in regulating learning rate. These results reveal a novel neural mechanism for setting learning rates and highlight acetylcholine’s role in adaptive learning.

Although the accumbal cholinergic system has long been known to play a crucial role in learning, its precise computational role has remained unspecified. Our results suggest that cue-evoked acetylcholine dips encode the cue’s associative salience, which is acquired during learning. This proposal explains not only our recording results but also previous findings in which acetylcholine activity was manipulated. For instance, mice with a selective knockout of the vesicular acetylcholine transporter (VAChTcKO) fail to discriminate between cues paired with reward (CS+) and non-reward (CS−) during acquisition of an appetitive association ^22^. In that study, acetylcholine dips to the CS+, observed in control mice, were absent in VAChTcKO mice^22^. Our findings suggest that without these dips, VAChTcKO mice could not assign appropriate salience to the cues, resulting in slower learning and a failure to learn the reward contingencies. Our proposal is also in line with several recent findings, explaining why tonic stimulation of cholinergic interneurons impairs reversal learning^46^, why learning is facilitated when acetylcholine dips precede dopamine RPEs^18^, and why preventing morphine-induced dips in acetylcholine also prevents morphine-induced associative learning^17^.

This proposal contrasts with previous studies that have concluded that acetylcholine signals in the dorsal striatum^47^ and basal forebrain^48^ encode RPEs. While the acetylcholine responses in NAcc are also selective to cue value and decrease in response to reward outcomes as learning progresses, similar to an RPE signal, we believe that acetylcholine in the NAcc does not directly encode RPEs as in the basal forebrain. First, during the initial learning phase, acetylcholine responses are not modulated by cues. With continued training, however, acetylcholine becomes progressively inhibited by both Go and No-Go cues, even though these two cues are associated with opposing RPE signals. Second, dopamine signals, known to encode RPEs, develop cue selectivity earlier than acetylcholine. Especially in the reversal learning task, the acetylcholine responses could not differentiate cues even when rats learned the reversed reward contingency without further training. These observations suggest that acetylcholine selectivity may be tied to more computationally complex variables like acquired salience, rather than directly encoding RPE or unexpected value. In contrast to our findings, Gallo et al.^49^ found that the D2R upregulation in nucleus accumbens cholinergic interneurons enhanced the cholinergic pause but delayed No-Go learning. This discrepancy may reflect differences in task structure, as their No-Go cue was itself associated with reward and signaled by a distinct visual cue, which may increase cue salience and encourage sign-tracking, potentially confounding the behavioral readout.Importantly, the correspondence between cue-evoked acetylcholine dips and the concept of associative salience does not rely on the specific form of the hybrid model used in this study. A similar model proposed by Esber and Haselgrove (2011)^34^, which also integrates PDS and UDS via a very different underlying theoretical mechanism, produces similar predictions. Unlike the Mackintosh and Pearce-Hall models, this model defines PDS as the sum of a cue’s association strength with positive and negative reinforcers, and UDS reflects the extent to which preceding events predict the cue itself. Total salience is the difference between these two components (Supplementary information). Nevertheless, when this model was applied to the Go/No-Go task, the PDS increased as cue-outcome associations were learned, and the UDS increased with repeated cue presentations. With an appropriate learning rate, total salience would initially increase and subsequently decline. Since the water reward supports stronger associations, the Go cue exhibited higher salience, consistent with predictions from our hybrid model (Extended Data Fig. 6). This robustness suggests that the salience carried by acetylcholine reflects a more general principle of learning, transcending the particular model adopted here.

The associative learning model we used in the current study proposes that acquired salience can modulate the learning rate and subsequent RPEs, while the RPEs are quantitatively related to the prediction uncertainty and can, in turn, affect salience. Such a mechanism highlights potential interactions between cholinergic and dopaminergic systems across trials. Supporting this, we observed significant effects of cue-evoked acetylcholine dips on the updates in dopamine and behavior in the current study. However, we failed to observe a reciprocal effect of the dopamine signal on the acetylcholine (Extended Data Fig. 3 B and C), possibly due to the complexity of salience dynamics, which may obscure the effects of individual factors. The neural mechanisms underlying this modulation remain unclear and require further investigation. Future work should also test the generalizability of our observations, especially in classical salience paradigms such as latent inhibition.

In summary, our findings revealed that associative learning is determined by a characteristic pattern of sequential dopamine and acetylcholine dynamics in the NAcc, and suggest that cue-evoked acetylcholine dips encode the associative or acquired salience of stimuli across various learning stages and tasks. We found surprising consistency between the acetylcholine dips and the salience term predicted by a hybrid attentional associative learning model, where associative salience was determined by both predictiveness-driven and uncertainty-driven salience. These findings, together with the observed causal effects of manipulation on accumbal cholinergic interneurons on behavior updates, support acetylcholine’s role in encoding salience and modulating learning rates.

## Figures

**Extended Data Fig. 1.**
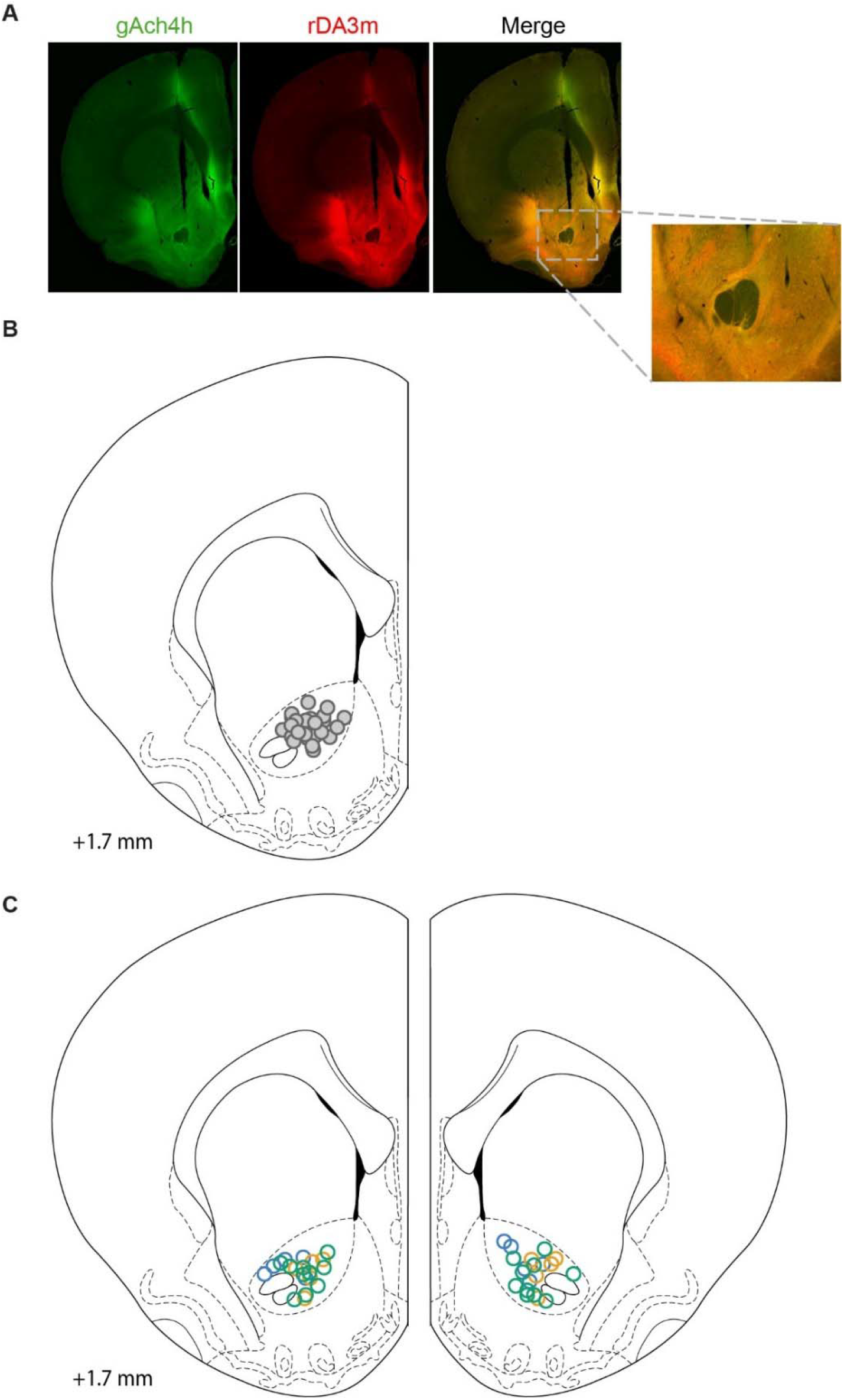
Histological verification and photometry recording locations. (**A**) Representative histological microphotograph with histological detection of both sensors. (**B**) Location of fiber tips in the NAcc for all recorded rats (N=22 rats). Among these, 5 rats had fibers implanted in both hemispheres and 18 rats had fibers implanted in a one hemisphere. (**C**) Location of fiber tips in the NAcc for all optogenetic rats (N = 20). All rats received bilateral fiber implants. Green, blue, and yellow dots indicate fiber-tip locations for the control (n = 10), activation (n = 5), and inhibition (n = 5) groups, respectively.

**Extended Data Fig. 2.**
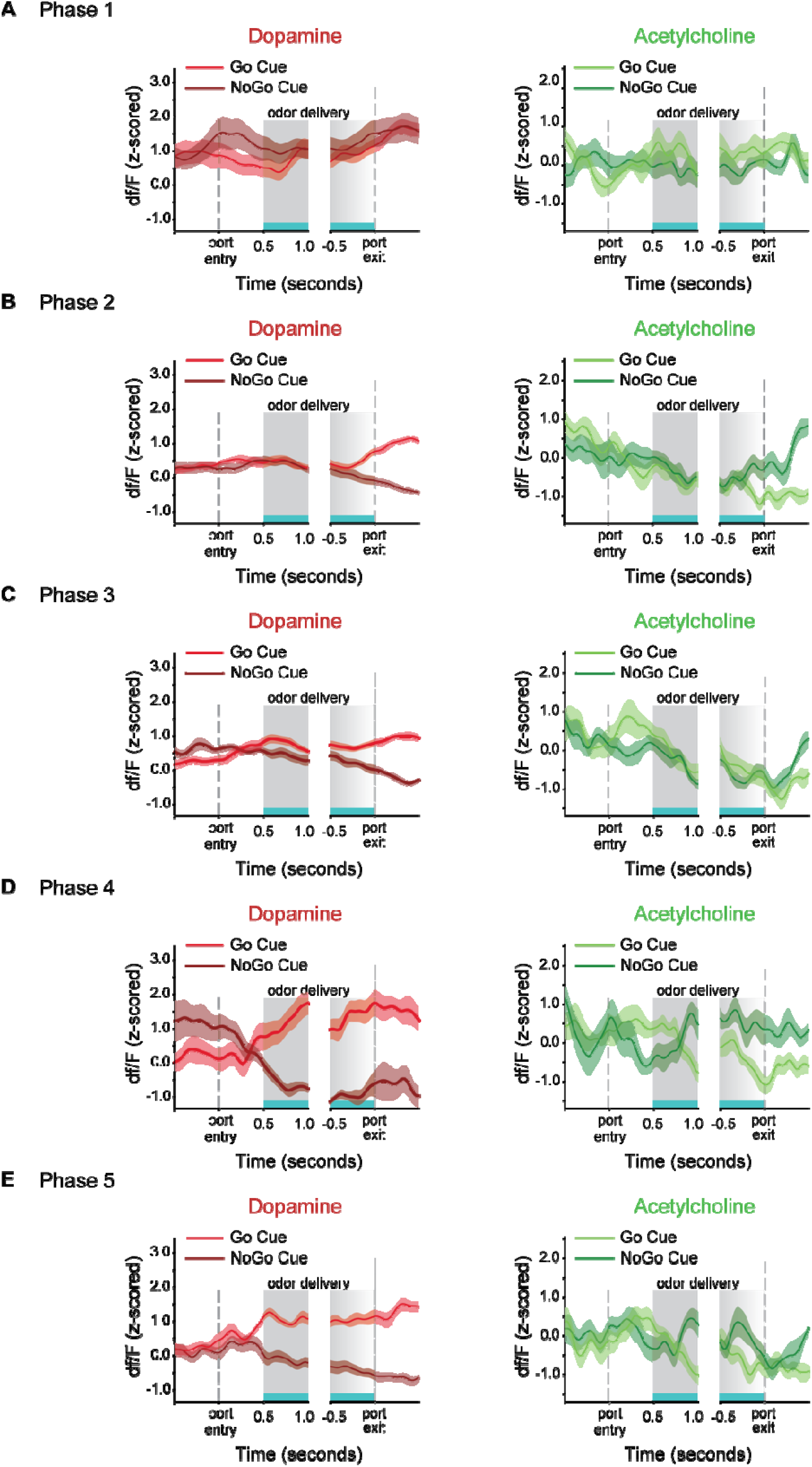
Representative acetylcholine and dopamine PSTHs from an example rat (Experiment 1) in different stages. (**A**) The first ten trials. (**B**) Ten trials immediately before and after dopamine selectivity emerged. (**C**) Ten trials immediately before and after behavioral criterion was met.

**Extended Data Fig. 3.**
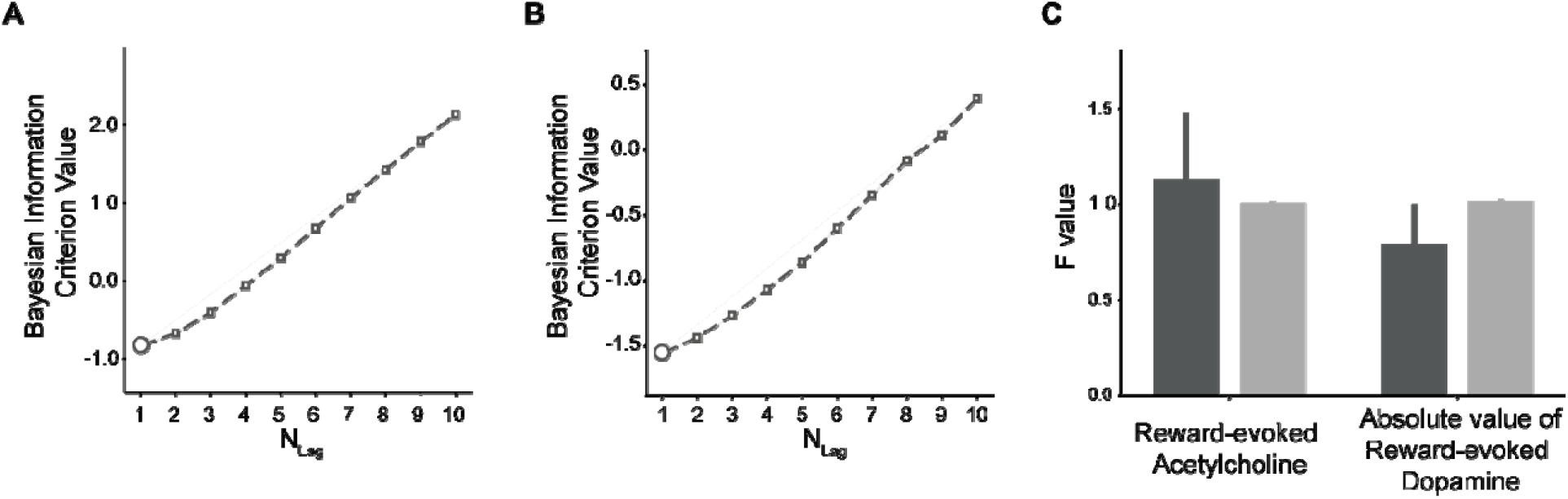
Determination of VAR lag length and Predict cue-evoked acetylcholine response with dopamine responses. (**A**) Bayesian Information Criterion (BIC) values for VAR models predicting cue value updates (fig. 3A) across different lag lengths. The circle indicates the lag (=1) with lowest BIC value. (**B**) BIC values for VAR models predicting cue-evoked acetylcholine signals based on reward-evoked dopamine signals, their absolute values, and past cue-evoked acetylcholine responses, with varying lag lengths. The circle indicates the lag (=1) with lowest BIC value. (**C**) F-values from the Granger causality test for the actual data (dark grey bars) versus shuffle data (light grey bars). No significant effects were found for reward-evoked dopamine signals (p-value=0.66) or their absolute value (p-value=0.45) on the subsequent cue-evoked acetylcholine signal. The error bars in indicate S.E across trials.

**Extended Data Fig. 4.**
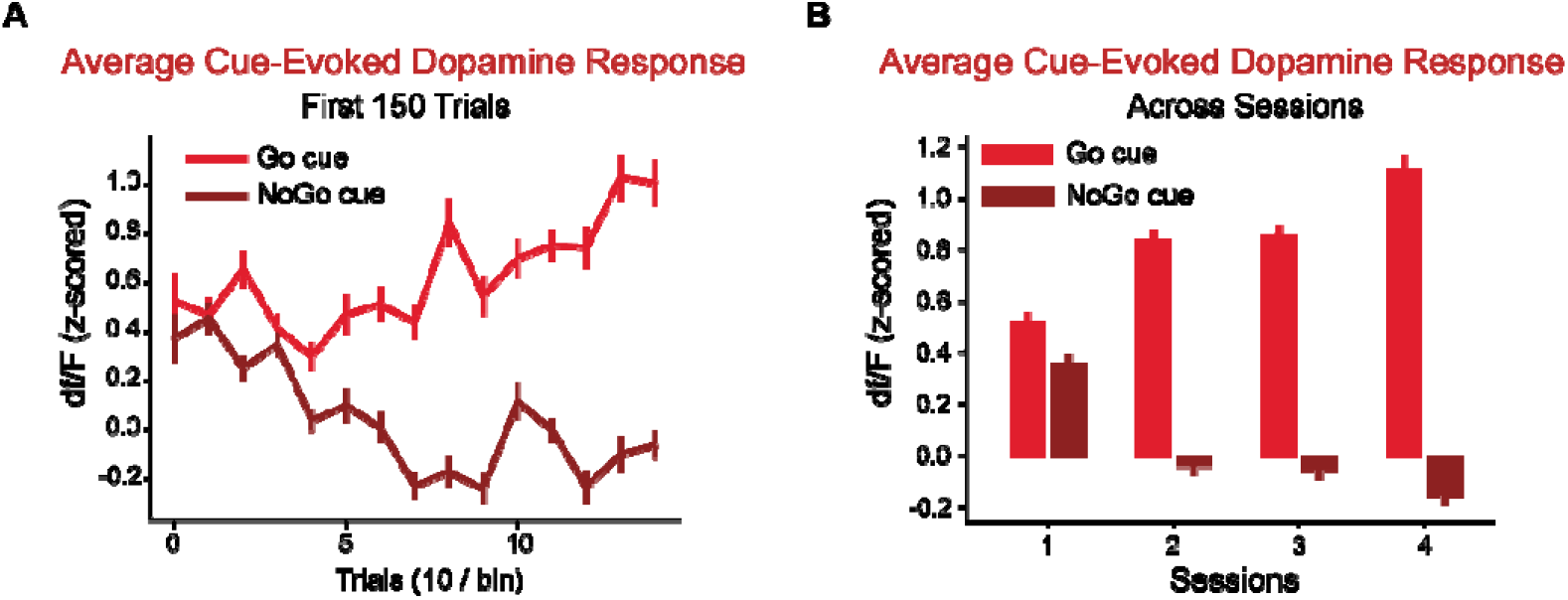
Dynamic changes of dopamine response across trials and sessions. (**A**) Average cue-evoked dopamine responses for Go (light red) and NoGo (dark red) cues in the first 150 trials. (**B**) Averaged cue-evoked dopamine responses for Go (light red) and NoGo (dark red) cues across training sessions. The error bars indicate S.E.M. across trials

**Extended Data Fig. 5.**
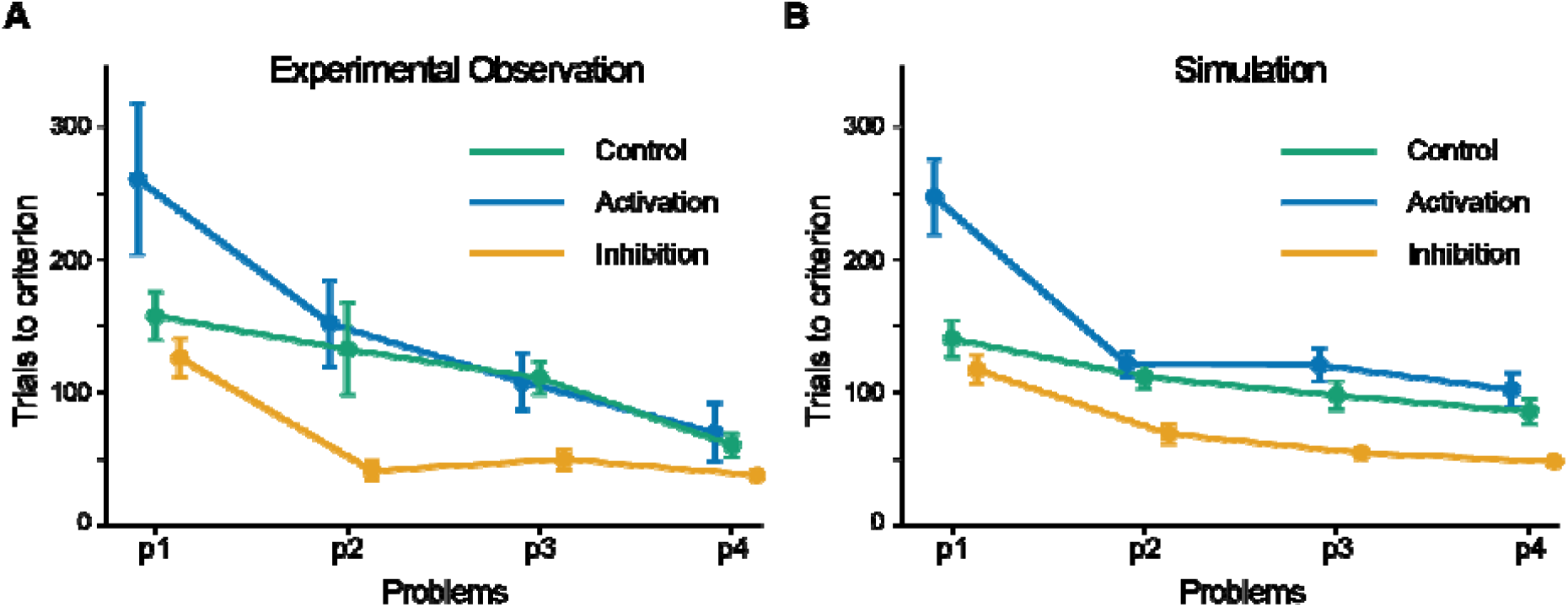
Observed and simulated effects of optogenetic manipulation across four learning problems. Trials to criterion across four sequential Go/No-Go problems in the experiment **(A)** and the extended hybrid model **(B)**. Control, activation, and inhibition conditions are shown in green, blue, and yellow, respectively. Error bars indicate the mean ± SEM across rats or simulations.

**Extended Data Fig. 6.**
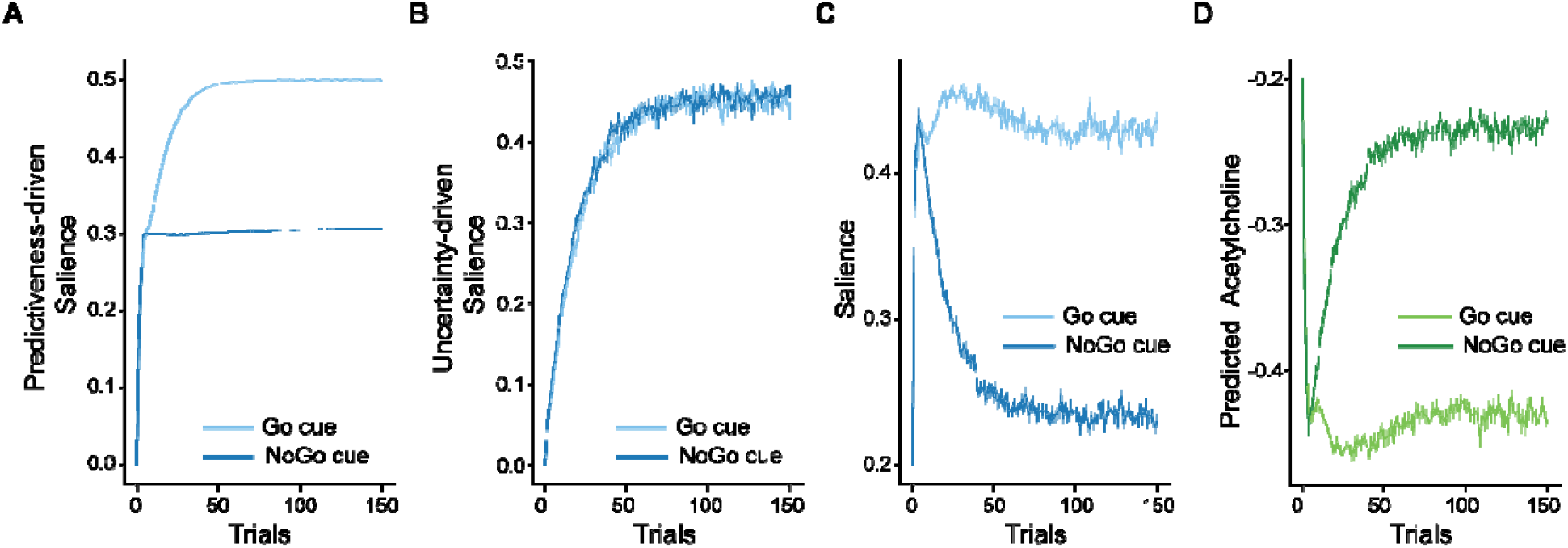
Esber and Haselgrove’s model explains the acetylcholine responses during the Go/NoGo task (Experiment 1). (**A**) The predictiveness-driven salience (PDS) component predicted by the E-H model. (**B**) The uncertainty-driven salience (UDS) component predicted by the E-H model. (**C**) The net salience calculated as the difference between PDS and UDS. (**D**) acetylcholine responses predicted by the E-H model. The error bars indicate S.E.M. across runs.

**Extended Data Fig. 7.**
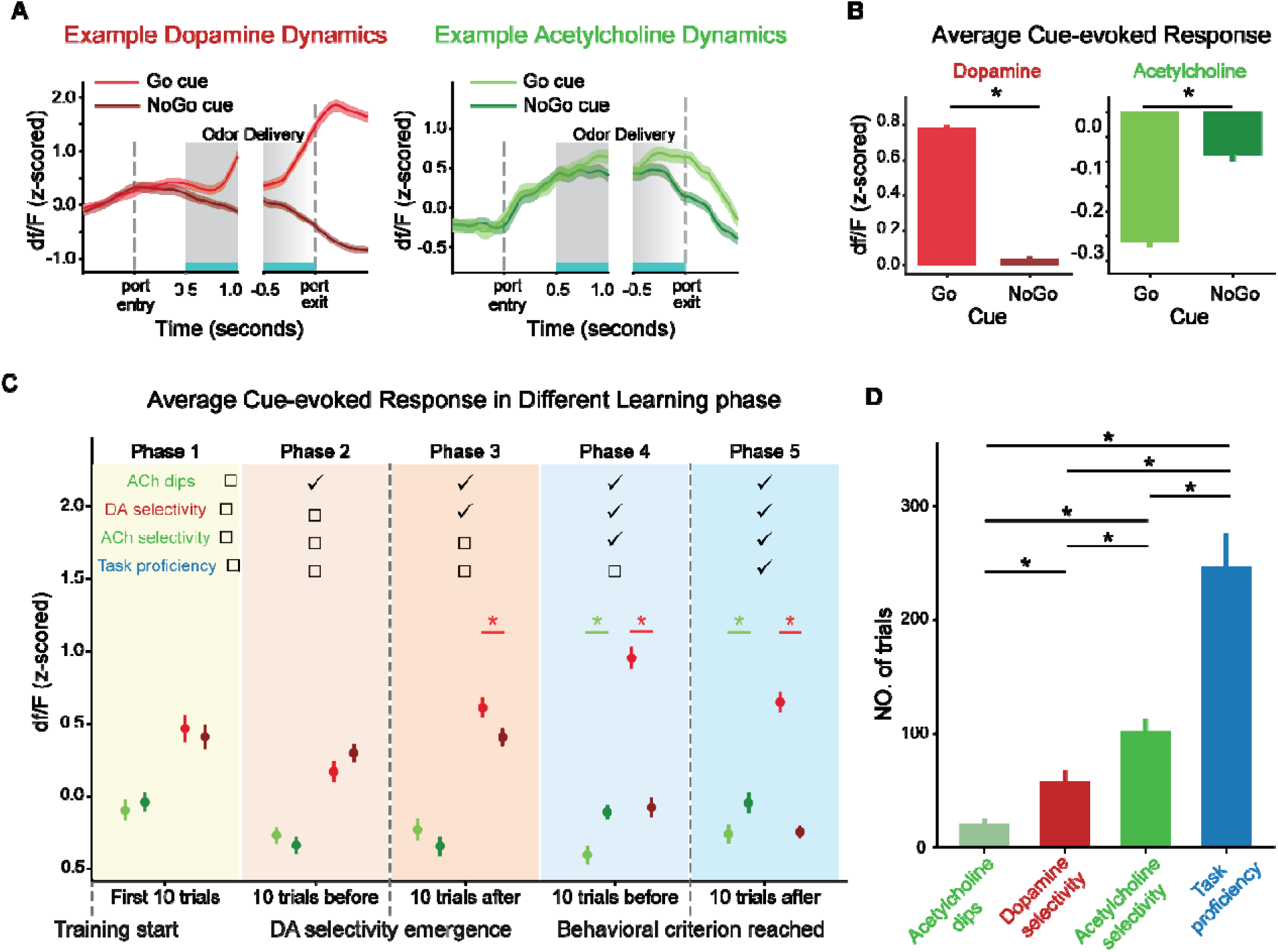
Dynamics of dopamine and acetylcholine during the Go/NoGo task (Experiment 1) including the outlier rat. **(A)**. The dopamine (left panel) and acetylcholine (right panel) dynamics within trials from the outlier rat. **(B-D)** Analyses as presented in fig. 2, now including the outlier rat.

## Methods

### Experimental model and subject details

We used 22 male Long–Evans rats weighing >450 g at the start of the experiment for fiber-photometry experiments and 20 ChAT-Cre rats for optogenetic experiments. The ChAT-Cre cohort included 12 males weighing >450 g and 8 females weighing >300 g at the start of the experiment. Rats were sourced from Charles River Laboratories and the NIDA IRP Breeding Facility. All rats were housed on a 12-h light-dark cycle at 25°C. They had free access to food but were water-restricted to approximately 85% of their original weight during the experiments. All rats received around 20 ml of water per day. Behavioral testing was performed during the light phase of the light-dark schedule. Experiment 1 was conducted on fourteen rats. One rat was excluded from the analyses due to markedly different acetylcholine signals, although including this rat did not alter the overall conclusions. The acetylcholine data for this rat, as well as the results for all fourteen rats combined, were provided in the supplementary materials (Extended Data Fig. 7). An additional eight rats were initially used in Experiment 3. One rat was not trained for Experiment 3 due to health issues, while two rats failed to meet the behavioral criterion in certain reversal problems and were therefore excluded from analyses. Five of the rats that were initially used in Experiment 1 were later involved in Experiment 4. For Experiment 2, each activation and inhibition group included three male and two female ChAT-Cre rats. The control group included six male and four female ChAT-Cre rats, divided evenly between the two light-delivery protocols. All procedures followed the National Institutes of Health guidelines, as determined by the National Institute on Drug Abuse Intramural Research Program (NIDA IRP) Animal Care and Use Committee (protocol no. 23-CNRB-108).

### Surgical procedures

#### Recording

Animals were anesthetized with isoflurane (3-5% for induction and 1-2% for maintenance). To simultaneously measure the dopamine and acetylcholine release in the nucleus accumbens core, we injected the D1 receptor-based red dopamine sensor-transfecting virus (pAAV-hsyn-rDA3m) and the muscarinic M3 receptor-based green acetylcholine sensor-transfecting virus (AAV-hSyn-gACh4h) into the unilateral NAcc of rats. The injection coordinates were AP +1.7 mm, ML ±1.7 mm, and DV -6.3 and -6.2 mm from the brain surface. Rats injected in the left and right hemispheres were counterbalanced. In the five rats used for both Experiment 1 and the long-term reversal learning task, the dopamine and acetylcholine sensor-transfecting viruses were injected into different hemispheres. For all other rats, both sensors were injected into the same hemisphere. A total of 1.0 μL of dopamine and acetylcholine sensor-transfecting virus, 0.5 μL for each, was delivered in each site at 0.1 μL/min via an infusion pump. All viruses were obtained from BrainVTA. The injection needle was held in for an additional 5 min after the completion of the injection to avoid the backflow of viruses. An optic fiber (0.37 NA, 200 μm diameter, Neurophotometrics, CA) was implanted in the most dorsal viral infusion location. Exposed fiber ferrules and a protective black 3D-printed headcap were secured to the skull with dental cement. After surgery, rats were given Cephalexin (15 mg/kg orally, once daily) for two weeks to prevent infections. Training started at least four weeks after the surgery to allow viral expression.

#### Manipulation

Optogenetic manipulation surgeries followed the same procedures as the recording surgeries, with only two differences. First, Cre-dependent viral vectors were injected bilaterally into the NAcc core: activation rats received pAAV-Ef1a-DIO-hChR2-EYFP, inhibition rats received pAAV-Ef1a-DIO-eNpHR-EYFP, and control rats received pAAV-Ef1a-DIO-EYFP. Second, bilateral PlexBright fiber-stub implants were placed above the dorsal-most viral infusion sites for light delivery. All other surgical procedures were identical to those used for the recording experiments.

### Fiber Photometry

#### Recording

Fluorescence signals were recorded using custom-ordered multi-pronged fiber optic patch cables (200 μm diameter, 0.37 NA, Doric Lenses, Canada) that were attached to the optic fiber ferrules on the skull of the rats with brass sleeves (Thorlabs, NJ). Up to 2 fibers were connected at a time in each rat for recordings, and they were shielded and secured with a custom 3D-printed headcap-swivel shielding system that allowed for the relatively free movement of the rats without the use of optic commutators and prevented the spillover of light during recordings.

Recordings were conducted using an FP3002 system (Neurophotometrics, CA), by providing fiber-coupled LEDs at 470 nm (green, active acetylcholine-dependent signal), 560 nm (red, active dopamine-dependent signal) and 415 nm (isosbestic reference signal) excitation light through the patch cord in interleaved LED pulses at 150 Hz (50 Hz acquisition rate for each channel). The light was reflected through a dichroic mirror and onto a 20×Olympus objective. Excitation power was measured at ∼150-200 µW at the tip of the patch cord. Emitted fluorescent light was captured through the same cords, separated with an image splitting filter, and captured via a high quantum efficiency CMOS camera. Signals were acquired and synchronized with behavioral events using Bonsai.

#### Pre-processing

We filtered the raw fluorescence signals from the 470 nm (active), 560 nm (active), and 415 nm (reference) channels by using a causal median filter and a second-order Butterworth low-pass filter with a cutoff frequency of 5 Hz. Next, we filtered each channel’s data with a second-order Butterworth high-pass filter with a cutoff frequency of 0.002 Hz, aiming to eliminate the exponential decay in fluorescence caused by factors like photobleaching. After that, the reference channel data were then fitted to each active signal using second-order polynomial regressions, and the fitted data were subsequently subtracted from the active channel and divided by the exponential fit of the active channel to remove signal-independent variations in fluorescence. Finally, the resulting active signal was z-scored for each session.

### Optogenetic manipulation

Light was delivered through PlexBright optical patch cables combined with PlexBright Compact LED Modules. For activation, 465 nm blue light (6–10 mW output) was delivered to activate hChR2-expressing accumbal cholinergic interneurons and increase the acetylcholine release. For inhibition, 590 nm yellow light (6–10 mW output) was delivered to activate eNpHR-expressing interneurons and decrease acetylcholine release. Control rats received the same 465 or 590 nm light-delivery protocols after injection of the control vector lacking opsin. During the Go/No-Go task, light was delivered continuously from 100 ms before odor onset until odor offset.

### Behavioral task

Rats were trained and recorded at least 4 weeks after the surgeries in standard operant boxes, which were aluminum chambers approximately 18 inches in height, depth, and width. A central odor port was located above the two fluid wells located on the left wall of the chamber, and one well was physically blocked and not used during this study. Two lights were located above the odor port. The odor port was connected to an airflow dilution olfactometer to allow the rapid delivery and removal of odor cues. Odors were chosen from compounds obtained from International Flavors and Fragrances (New York, NY). Odor delivery and fluid delivery to the odor port and fluid well were controlled by the behavioral computer via a system of flow meters and solenoids. The entry into the odor port and fluid well was detected by the interruptions of an infrared beam, which was connected to the behavioral computer.

All experiments followed the same procedures in a single trial and differed only in the odor used and reward contingency. After the illumination of lights inside the box, rats were allowed to perform a nosepoke into the central odor port to initiate a new trial. Following a 500 ms delay, an odor cue was presented for at least 500 ms. The presented odor was randomly selected from two possible cues that differed in different experiments and problems, and determined the reward availability in the fluid well. After the initial 500 ms of odor delivery, the rats were free to leave the odor port and odor delivery was terminated upon port exit. Rats then had 3 seconds to make a response at the fluid well. If a response was initiated, 0.05 ml water was delivered 500 ms later in reward trials, while no fluid was delivered in non-reward trials. During reward trials, the panel lights remained illuminated until the rat exited the fluid well, and the lights turning off indicated the end of a trial. In non-reward trials, the lights were turned off immediately after the response. If the rats did not respond within 3 seconds after leaving the odor port, the lights were extinguished then. Inter-trial intervals commenced upon the turnoff of the panel lights and lasted for 4 seconds, extending to 8 seconds following responses in non-rewarded trials. Before involving any experiment, rats were trained with no odor and received water rewards at the fluid well to familiarize themselves with the task procedures.

#### Experiment 1 (Go/No-Go task)

Fourteen naive Long–Evans male rats were included in Experiment 1 and trained with a Go/No-Go task using the task structure mentioned above. Two distinct odor cues were used: one “Go” cue associated with a reward and the other “No-Go” cue associated with no reward (Fig. 1B). To keep an overall balance in the number of positive vs negative trials throughout the session, we included 25 positive and 25 negative trials in every 50 trial block, but the order was pseudo-randomly generated independently. Rats were trained to learn the associations between cues and outcomes for at least six sessions.

#### Experiment 2 (four sequential Go/No-Go tasks)

Twelve male ChAT-Cre rats and eight female ChAT-Cre rats were trained on a sequence of four Go/No-Go problems, each using a unique odor pair (Fig. 4B). For each problem, training continued until 20 trials after rats reached the behavioral criterion. Rats then advanced to the next problem in the following session, continuing until they had acquired all four problems.

#### Experiment 3 (reversal task)

Seven naive Long–Evans male rats were trained with three Go/No-Go problems and were then involved in Experiment 3. Data from two were excluded from analyses since they failed to meet the behavioral criterion in certain reversal problems. Experiment 3 consisted of three reversal problems (Fig. 5A), each involving two stages. In the first stage, the rats were trained with a Go/No-Go problem used in Experiment 1. Once rats achieved the behavioral criterion on the initial problem, 18 correct responses within a 20-trial moving block, training was stopped 20 trials later to ensure that rats were equally proficient. The second stage was initiated once rats reached the criterion again in the next session. In the second stage, the reward contingencies between cues and outcomes were reversed, i.e., the cue previously associated with a reward now indicated no reward, and the cue previously associated with no reward led to a reward. To encourage rats to learn this reversal, we introduced three teaching trials after each reversal, in which the previously unrewarded cues were presented, and rats had unlimited time to respond at the fluid well. These teaching trials increased the probability that rats responded at the fluid well and discovered that the reward contingency was reversed. Training with reversed contingencies continued over multiple sessions until the rats met the same behavioral criterion. After the successful completion of the first reversal, the reward contingencies for the second and third sets of odor pairs were similarly trained and subsequently reversed.

#### Experiment 4 (reversal task with extensive training)

Five Long–Evans male rats used in Experiment 1 were trained with a Go/No-Go task over six sessions. In the seventh session, we reversed the reward contingencies 20 trials after the rats achieved the behavioral criterion. Training with the reversed contingencies then continued for ten sessions.

### Histological procedures

After completion of the experiment, rats were perfused with chilled phosphate buffer saline (PBS) followed by 4% paraformaldehyde in PBS. The brains were then immersed in 20% sucrose in PBS for at least 24 hours and frozen. The brains were sliced at 50 μm in series for the determination of optic fiber implant location in the NAcc core and were then used for fluorescent immunohistochemistry.

Sections from rats used for fiber-photometry recordings were further stained with DAPI (Vectashield-DAPI, Vector Lab, Burlingame, CA), and processed for immunohistochemical detection of green and red fluorescent protein. For fluorescence immunohistochemistry, gACh4h was immunostained using a chicken anti-GFP antibody (1:1,000, Abcam, catalog #ab13970) in 0.1% Triton X-100/1× PBS and incubated in Alexa Fluor 488 AffiniPure Donkey Anti-Chicken IgY (IgG) secondary antibody (H+L, code: 703-545-155, RRID: AB_2340375, dilution: 1:100) overnight. rDA3m was immunostained using a rabbit anti-RFP antibody (1:1,000, Rockland, catalog #600-401-379) in 0.1% Triton X-100/1× PBS and followed by incubation in Alexa Fluor 594 AffiniPure Donkey Anti-Rabbit IgG secondary antibody (H+L, code: 711-585-152, RRID: AB_2340621, dilution 1:100). Fluorescence microscopy images were acquired using a BZ-X800 microscope (Keyence). Fluorescent expression patterns and optic fiber locations were identified from the acquired images and superimposed onto corresponding anatomical atlas templates.

For rats used in optogenetic experiments, free-floating coronal sections were rinsed three times for 10 min each in phosphate buffer and incubated for 1 hour at room temperature in blocking solution containing 4% bovine serum albumin and 0.3% Triton X-100 in phosphate buffer. Sections were then incubated overnight at 4°C with a mouse anti-GFP primary antibody (1:500; Takara Bio, catalog no. 632381) and rabbit anti-VAChT (1:500; Frontier Institute Co., Ltd., Japan, catalog no. MSFR106490) primary antibodies. The next day, sections were rinsed three times for 10 min each in phosphate buffer and incubated for 2 hours with an Alexa Fluor 488-conjugated secondary antibody and Alexa Fluor 594-conjugated anti-rabbit secondary antibodies (1:200; Jackson ImmunoResearch Laboratories). After washing, sections were mounted onto glass slides. Fluorescence images of sections were acquired using an Olympus VS200 slide scanner. Low-magnification images were obtained using a 10× objective, acquiring fluorescence channels to minimize spectral overlap.

### Detection of acetylcholine dips and neuromodulatory selectivity

Cue-evoked dopamine and acetylcholine responses were quantified for each completed trial as the mean z-scored fluorescence from odor onset to odor-port exit. Within each rat, Go and NoGo trials were ordered separately according to their cue-specific trial ordinal and smoothed using a causal half-Gaussian kernel (θ = 0.5). Kernel weights decreased with increasing lag, assigning the greatest weight to the current trial while excluding all future trials.

To estimate response onset, trailing windows containing 10 Go and 10 NoGo trials were evaluated at successive cue-ordinal positions. ACh dip onset was defined as the first position at which the combined Go and NoGo ACh responses differed significantly from a zero-valued vector. DA-selectivity and ACh-selectivity onsets were defined as the first positions at which Go and NoGo responses differed significantly for the corresponding neuromodulator. All onset tests were performed using two-sided, equal-variance *t*-tests with a significance threshold of *p* <0.01. Because Go and NoGo cues occurred at approximately equal frequencies, the detected cue-ordinal position was multiplied by two to estimate total completed-trial counts. Pairwise differences among ACh dip onset, DA-selectivity onset, ACh-selectivity onset, and behavioral criterion were assessed at the rat level using the paired Bayesian bootstrap procedure described below.

### Vector autoregression modeling and Granger causality testing

The hypothesis that the level of acetylcholine release evoked by a cue modulates the learning rate led to the prediction that the acetylcholine release, in combination with the prediction error induced by the reward outcome, could significantly influence the updating of the cue’s value. Prediction error evoked by the reward outcome can be estimated by the corresponding dopamine response, while the cue’s value updates can be inferred from changes in the cue-evoked dopamine response across trials.

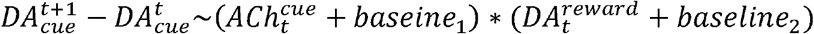

To test this relationship, we applied a vector autoregression model separately to all trials for each condition separately. This analysis assessed the correlation between the cue-evoked dopamine response, the cue-evoked acetylcholine response, the reward-evoked dopamine response, and their interaction term. The model is expressed as:

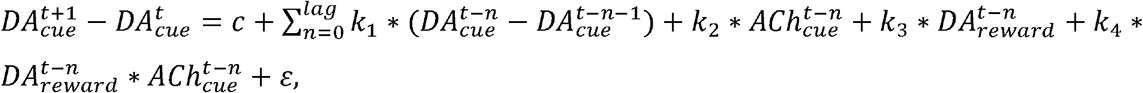

where *c* is a constant, *k*_*i*_ are model parameters and *ε* is the error term. Bayesian information criterion (BIC) was calculated for different lags, and the lag with the lowest information criterion value was selected for further analysis (Extended Data Fig. 3a).

Next, the model was fitted separately for each session and trial type (presented odor). Granger causality analysis was then applied to determine whether the cue-evoked acetylcholine response, reward-evoked dopamine response, and their interaction term provided significant information about the cue-evoked dopamine response beyond what could be explained by its past values. The F values were from the original data compared to those obtained from the control data, where the were shuffled 1000 times, to determine the statistical significance of each factor’s effect on cue-evoked dopamine responses.

### Bayesian bootstrap analysis

Bayesian bootstrap mean-difference tests were used for comparisons of both the temporal emergence of neural and behavioral measures and the effects of optogenetic manipulation on learning. Comparisons among acetylcholine dip onset, dopamine selectivity onset, acetylcholine selectivity onset, and the behavioral criterion were performed using a paired Bayesian bootstrap, in which the same Dirichlet-distributed weights were applied jointly to all measures from each rat to preserve their within-animal correspondence. For comparisons between optogenetic groups, Dirichlet weights were sampled independently within each group. Bayesian bootstrap estimated uncertainty in group means and pairwise differences without relying on normality or equal-variance assumptions, which was suitable for the relatively small sample sizes in these analyses.

Learning performance was quantified as the number of trials required to reach the behavioral criterion. Post-criterion performance was quantified as the proportion of correct No-Go trials during the 20 trials after criterion was reached. Based on the hypothesis that reducing accumbal cholinergic interneuron activity facilitates learning, whereas increasing activity slows learning, planned directional comparisons predicted that trials to criterion would follow activation > control > inhibition, whereas post-criterion No-Go performance would follow inhibition > control > activation.

For pairwise group comparisons, behavioral metrics were first averaged across problems within each rat to reduce problem-level variability. For each planned comparison, observations from the two groups were treated as independent empirical distributions. *x*_*g,i*_ denotes the behavioral measure for animal (i) in group (g). For each bootstrap draw (b), independent Dirichlet-distributed weights were sampled within each group, and weighted group means were calculated as:

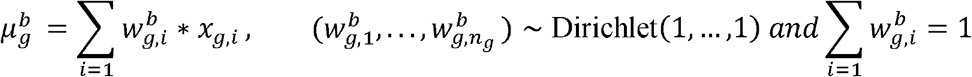

The bootstrapped mean difference for each draw was then computed as:

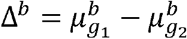

The posterior distribution of the mean difference was obtained from the distribution of Δ^*b*^ across bootstrap draws. For each planned directional comparison, the expected direction was encoded as

*s* = +1 when the prediction was Δ^*b*^ < 0, and *s* = −1 when the prediction was Δ < 0. The one-sided Bayesian-bootstrap tail probability was calculated as the posterior mass in the non-predicted direction:

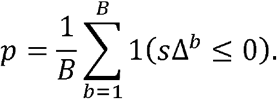

These values are reported as Bayesian-bootstrap posterior tail probabilities. When no bootstrap samples fell in the non-predicted direction, the tail probability was reported as *p* < 1/*B*. With B = 10,000 bootstrap draws, this corresponded to *p* < 1.0*e* −4.

### Attentional associative learning models

We used three attentional associative learning models to simulate the salience dynamics of stimuli in the learning process. The mechanisms for updating associative strength were the same across models, differing only in how salience was defined. By varying the definition of salience, three models provide different predictions on the salience dynamics during learning.

The associative strength for a conditioned stimulus, *A*, was represented by an excitatory component, *v*_*A*_, and an inhibitory component, 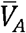, indicating the expectation of positive (water reward) and negative reinforcer (timeout), respectively. The net associative strength, 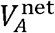, is calculated as:

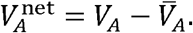

The updates to these components are driven by the prediction error, *δ*_*v*_, defined as:

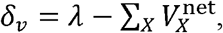

where *X* represents the set of stimuli represented in the environment and *λ* is the signed maximum associative strength supported by the reinforcer, representing either the reward or penalty value.

Specifically, λ is set to 1 for water reward, -0.2 for reward omission, and -0.25 for timeout penalty. The models update the excitatory and inhibitory components of the associative strength based on the sign and magnitude of the *δ* _*v*_,

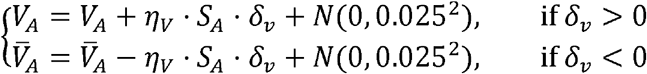

where *η*_*V*_ is the learning rate for associative strength, *S*_*A*_, indicating the salience of stimulus *A*. Salience is constrained to the interval, [0.01,1], to maintain a nonzero contribution to learning and modulates the influence of the prediction error on changes in associative strength. *S*_*A*_ modulates the overall impact of the *δ*_*v*_ on the associative strength and *N* (0,0.025) is the Gaussian noise with mean 0 and variance 0.025.

#### The predictiveness model

The predictiveness model is grounded in the Mackintosh model. The salience, *S*, is determined by the predictiveness, *α*, of a stimulus. Stimuli providing higher predictiveness compared to other perceivable stimuli are assigned higher salience. The change in *α*, δ _*α*_, is defined as the difference between the absolute error for other stimuli (*X* ≠ *A*) and the absolute error for the target stimulus (*A*):

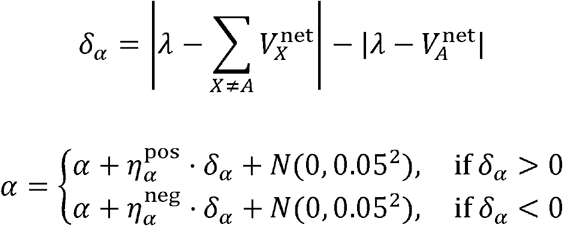

where 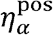 and 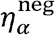 are the learning rates for increasing and decreasing *α*, respectively. To ensure that *α* remains within a meaningful range, it is rectified according to an upper limit, *α*_upper_, and a lower limit, *α*_lower_. *α*_lower_ is pre-defined, while *α*_upper_ is dynamic and determined by the absolute strength of the reinforcer.

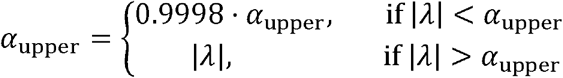

#### The uncertainty model

The uncertainty model is grounded in the Pearce-Hall model. The salience of a stimulus is determined by its uncertainty. Stimuli, which provide lower predictiveness and indicate higher uncertainty about the upcoming observations, are assigned higher salience, *σ*. The error in *σ,δ*_*σ*_, is defined as:

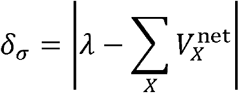

Then is updated similarly to *σ*, and the learning rate varies based on whether *δ*_*σ*_ is larger or smaller than the previous *σ*:

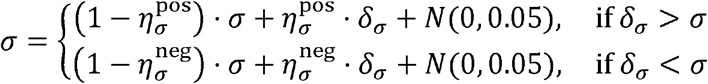

#### The hybrid model

The hybrid model integrates salience components from both the Mackintosh and Pearce-Hall models. The salience, *S*, is determined by the weighted sum of predictiveness-driven and uncertainty-driven salience of a stimulus.

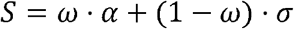

where 0< *ω* <1 determines the relative contribution of *α* and *σ*. In all simulations, *ω* is set to 0.3. The dynamics and update rules of *α* and *σ* are the same as those described in the predictiveness model and the uncertainty model.

The total salience of each cue is constrained to [0.01, 1] in all three models to prevent implausibly low or high values.

#### Go/No-Go task

We simplified the task structure in our simulations. Completed trials from rats were used to train the models. For each trial, we assumed that two stimuli were represented: the presented odor and the background stimuli in the environment. Following these cues, one of three reinforcers was presented: water reward, reward omission, and timeout penalty. In all three models, 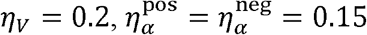, and 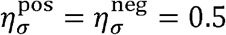.

To evaluate the goodness of fit for the acetylcholine signal in the Go/No-Go task, we used rat behavioral data and fitted models by minimizing the mean squared error in reproducing acetylcholine responses. This optimization was performed using differential evolution. Due to the simulation’s complexity, we fitted three key free parameters in each model that we believe most significantly affected the results. For the predictiveness model, we relaxed the assumption that 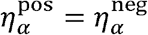 and set 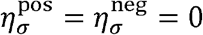. The fitted parameters were, 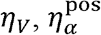 and 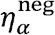. For the uncertainty model, we relaxed the assumption that 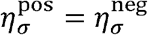, and set 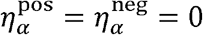. The fitted parameters were 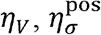 and 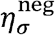. For the hybrid model, we assumed 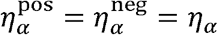 and 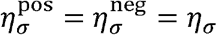. The fitted parameters were *η*_*V*_, *η* _*α*_ and *η* _*σ*_. We compared the losses from all entries in the population generated by the fitting algorithm, i.e., differential evolution, across different models to evaluate their relative fitting performance.

#### Optogenetic Manipulations

To determine whether the hybrid model could account for the behavioral effects of optogenetic manipulation, we simulated Go/No-Go acquisition under three conditions. Optogenetic activation was modeled by decreasing the trial-by-trial salience generated by the hybrid model by 0.2, whereas optogenetic inhibition was modeled by increasing salience by 0.2. Salience was unchanged in the control condition. In all simulations, salience was constrained to the same range used in the other simulations, [0.01, 1]. All other model parameters were held constant across conditions.

On each trial, the model selected between Go and No-Go actions according to:

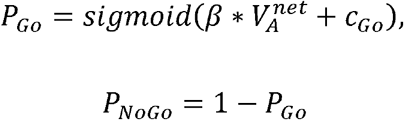

where *β* = 50 controls the sensitivity of choice to associative strength, and *c*_*Go*_ = 0 15 represents a value-independent bias toward making a Go response. Actions were sampled probabilistically from the resulting choice probabilities.

For each condition, 20 independent simulations were performed. The behavioral criterion was defined as the first trial at which performance reached at least 90% accuracy in the last 20-trial window, matching the criterion used to analyze the behavioral data.

#### Reversal task

To capture the dynamics of the acetylcholine signal in the reversal task, we relaxed the assumptions 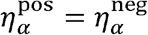 and 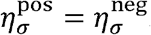. The results shown in Fig. 7 were obtained using the parameters: 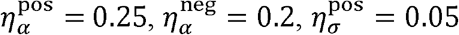, and 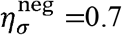. All other parameters were consistent with those used in the Go/No-Go task.

#### The extended hybrid associative learning models explain the problem-dependent effects of optogenetic manipulation.]

To test whether the behavioral changes observed across Go/No-Go problems during optogenetic manipulation could be captured by the associative learning model, we extend the hybrid model with two mechanisms that allow knowledge acquired from previous problems to facilitate learning in new ones.

First, the salience baseline of odor cues, *S*_*b,p*_, shared by all odor cues is updated according to the salience assigned to odors in the previous problem, reflecting the generalization of learned cue-outcome associations across problems. As a result, the salience baseline increases over problems, reducing the number of trials required to reach the behavioral criterion in later problems.

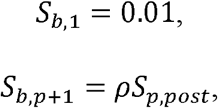

where *ρ* = 0.65, and *S*_*p,post*_ is the mean salience of the two odor cues over the 20 trials from the previous problem. The resulting baseline remains fixed throughout the next problem.

As in the main text, optogenetic manipulation alters the associative salience of the presented cue. Activation and inhibition are modeled as additive offsets of −0.25 and +0.7, respectively. After applying these offsets, salience is bounded below by the salience baseline and above by 1. Optogenetic manipulations do not directly change the salience baseline, but their effects on learning could indirectly influence the salience baseline transferred to the next problem.

Second, we included an experience-dependent Go bias. Before the Go/No-Go tasks, rats are trained on a shaping task without odor cues to learn the general task structure, in which all trials are rewarded. This experience produced a strong initial tendency for rats to approach the fluid well, which is captured by this Go bias term in the model. This bias decreases as rats experience trials in which approaching the well results in no reward but a timeout penalty.

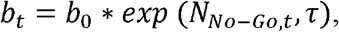

where *b*_0_ = 15, *τ*= 250, and N_*N o*−*Go,t*_ is the number of No-Go cue presentations before the trial *t*.

All other settings and parameters are the same as used in the main text. With these two mechanisms, the hybrid model captures the faster learning across problems and accounts for the observed problem-dependent effects of optogenetic manipulation (Extended Data Fig. 5). Intuitively, in the first problem, the salience baseline is close to zero, and optogenetic activation effectively reduces salience and therefore strongly disrupts learning. In later problems, the salience baseline enhanced by prior experience limits how far activation can reduce the salience. In contrast, inhibition increases salience above the baseline across all problems. However, in the first problem, the strong initial Go bias limits how quickly associative learning translates into No-Go behavior. Thus, inhibition accelerates learning but fails to produce a comparable behavioral improvement in the first problem. As the bias declines across problems, faster learning translates more readily into correct No-Go responses, accounting for the stronger behavioral effects of inhibition in later problems.*The attentional associative learning models based on Esber and Haselgrove (2011)*

Esber and Haselgrove (2011) is another influential hybrid attentional associative learning model (hereafter referred to as the E-H model). Here, we adopted the E-H model to reproduce the acetylcholine responses we observed in the Go/No-Go task. The Go/No-Go task structure was formalized as we did for the other models in the main text.

Like the models discussed in the main text, the E-H model represents a conditioned stimulus’s associative strength through two components: an excitatory component for positive reinforcer associations and an inhibitory component for negative reinforcer associations. The associative updating mechanism follows similar principles to the previously discussed models, a distinction lies in its updating pattern. The E-H model updates both positive and negative associations on every trial, rather than selectively updating positive associations only during trials with positive RPE and negative associations only during trials with negative RPE.

Similar to Model 3, the E-H model incorporates both predictiveness-driven and uncertainty-driven components in calculating salience. However, these two salience components are defined differently from previous models. In the E-H model, the predictive-drive salience, *α*, is defined as the sum of positive and negative association strength and we updated *α* in a temporal difference manner. This follows from the assumption that the cue will signal emotionally potent positive and negative reinforcers simultaneously, which have additive effects on salience:

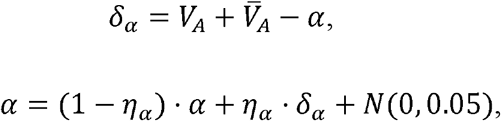

Unlike Le Pelly’s model, which focuses on outcome prediction uncertainty, the E-H model shifts the focus from outcome uncertainty to cue uncertainty, i.e., the uncertainty of the cue representation based on its context and preceding cues. Thus, the uncertainty-driven salience, *σ*, decreases as the cue becomes more predictable from contextual and preceding events.

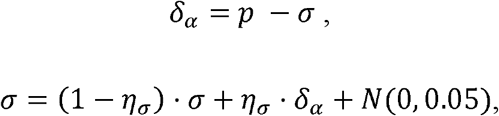

where *p*= 1 when cue A is represented in the current trial. Otherwise, *p* = 0.

The E-H model determines a cue’s overall salience, *S*, by calculating the difference between its predictiveness-driven and uncertainty-driven components, rather than their sum. This approach aligns with the model’s assumption that a cue becomes less salient when it becomes predictable.

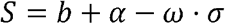

where *ω* = 0.6determines the relative contribution of *α* and *σ*

.*b* = 0.2 is the baseline for the salience. When we ran this model, *η* _*V*_ = 0.4, *η* _*α*_ = 0.5, and *η* _*σ*_ = 0.05.

## ACKNOWLEDGMENTS

We thank Nicolas Trisch for his constructive comments. We also thank Douglas Deutsch, Shiliang Zhang and the NIDA IRP Histology and Imaging Core for assistance with antibody testing and histological processing. This research was supported by the Intramural Research Program of the National Institutes of Health (NIH). The contributions of the NIH authors are considered Works of the United States Government. The findings and conclusions presented in this paper are those of the author(s) and do not necessarily reflect the views of the NIH or the U.S. Department of Health and Human Services.

## AUTHOR CONTRIBUTIONS

Z.Z., K.M.C., and G.S. conceived the study; Z.Z., K.M.C., W.Z., Y.Z., G.L., and Y.L. developed the methodology; Z.Z. and K.M.C. performed the investigation; Z.Z. and K.M.C. performed the visualization; G.S. supervised the study; Z.Z. wrote the original manuscript; and Z.Z., K.M.C., W.Z., G.S., Y.Z., G.L., and Y.L. reviewed and edited the manuscript.

## DATA AND METAERIALS AVAILABLITY

The data generated in this study will be deposited in the Zenodo database and accessible upon acceptance. All custom code used for reported analyses in this study will be available at GitHub upon acceptance.

## References

1 Costa, K. M. & Schoenbaum, G. Dopamine. Current Biology 32, R817–R824 (2022).

2 Cox, J. & Witten, I. B. Striatal circuits for reward learning and decision-making. Nature Reviews Neuroscience 20, 482–494 (2019).

3 Dudhabhate, B. B. & Costa, K. M. A brief history of dopamine prediction errors. Front Comput Neurosci 20, 1825622 (2026). 10.3389/fncom.2026.1825622

4 Costa, K. M. et al. Striatal dopamine signals errors in prediction across different informational domains. Sci Adv 11, eadq9684 (2025). 10.1126/sciadv.adq9684

5 Reynolds, J. N. et al. Coincidence of cholinergic pauses, dopaminergic activation and depolarisation of spiny projection neurons drives synaptic plasticity in the striatum. Nature Communications 13, 1296 (2022).

6 Shen, W. et al. M4 Muscarinic Receptor Signaling Ameliorates Striatal Plasticity Deficits in Models of L-DOPA-Induced Dyskinesia. Neuron 88, 762–773 (2015). 10.1016/j.neuron.2015.10.039

7 Picciotto, M. R., Higley, M. J. & Mineur, Y. S. Acetylcholine as a neuromodulator: cholinergic signaling shapes nervous system function and behavior. Neuron 76, 116–129 (2012).

8 Collins, A. L., Aitken, T. J., Greenfield, V. Y., Ostlund, S. B. & Wassum, K. M. Nucleus accumbens acetylcholine receptors modulate dopamine and motivation. Neuropsychopharmacology 41, 2830–2838 (2016).

9 Mohebi, A., Collins, V. L. & Berke, J. D. Accumbens cholinergic interneurons dynamically promote dopamine release and enable motivation. Elife 12, e85011 (2023).

10 Crespo, J. A., Stöckl, P., Zorn, K., Saria, A. & Zernig, G. Nucleus accumbens core acetylcholine is preferentially activated during acquisition of drug-vs food-reinforced behavior. Neuropsychopharmacology 33, 3213–3220 (2008).

11 Belilos, A. et al. Nucleus accumbens local circuit for cue-dependent aversive learning. Cell reports 42 (2023).

12 Aosaki, T. et al. Responses of tonically active neurons in the primate’s striatum undergo systematic changes during behavioral sensorimotor conditioning. J Neurosci 14, 3969–3984 (1994). 10.1523/JNEUROSCI.14-06-03969.1994

13 Yamada, H., Matsumoto, N. & Kimura, M. Tonically active neurons in the primate caudate nucleus and putamen differentially encode instructed motivational outcomes of action. Journal of Neuroscience 24, 3500–3510 (2004).

14 Costa, K. M. et al. Dopamine and acetylcholine correlations in the nucleus accumbens depend on behavioral task states. Current Biology 35, 1400-1407.e1403 (2025). 10.1016/j.cub.2025.01.064

15 Krok, A. C. et al. Intrinsic dopamine and acetylcholine dynamics in the striatum of mice. Nature 621, 543–549 (2023).

16 Chantranupong, L. et al. Dopamine and glutamate regulate striatal acetylcholine in decision-making. Nature 621, 577–585 (2023). 10.1038/s41586-023-06492-9

17 Yousefzadeh, S. A. et al. A cholinergic hub in the nucleus accumbens gates opioid-reward learning. Nature (2026). 10.1038/s41586-026-10887-9

18 Jang, H. J., McMahon Ward, R., Golden, C. E. M. & Constantinople, C. M. Acetylcholine demixes heterogeneous dopamine signals for learning and moving. Nat Neurosci 29, 840–850 (2026). 10.1038/s41593-026-02227-x

19 Collins, A. L. et al. Nucleus accumbens cholinergic interneurons oppose cue-motivated behavior. Biological psychiatry 86, 388–396 (2019).

20 Brown, M. T. et al. Ventral tegmental area GABA projections pause accumbal cholinergic interneurons to enhance associative learning. Nature 492, 452–456 (2012).

21 Duhne, M., Mohebi, A., Kim, K., Pelattini, L. & Berke, J. D. A mismatch between striatal cholinergic pauses and dopaminergic reward prediction errors. Proceedings of the National Academy of Sciences 121, e2410828121 (2024).

22 Skirzewski, M. et al. Continuous cholinergic-dopaminergic updating in the nucleus accumbens underlies approaches to reward-predicting cues. Nat Commun 13, 7924 (2022). 10.1038/s41467-022-35601-x

23 Le Pelley, M. E. The role of associative history in models of associative learning: A selective review and a hybrid model. Quarterly Journal of Experimental Psychology Section B 57, 193–243 (2004).

24 Le Pelley, M. & McLaren, I. Learned associability and associative change in human causal learning. The Quarterly Journal of Experimental Psychology: Section B 56, 68–79 (2003).

25 Griffiths, O. & Mitchell, C. J. Selective attention in human associative learning and recognition memory. Journal of Experimental Psychology: General 137, 626 (2008).

26 Kaye, H. & Pearce, J. M. The strength of the orienting response during Pavlovian conditioning. Journal of Experimental Psychology: Animal Behavior Processes 10, 90 (1984).

27 Hogarth, L., Dickinson, A., Austin, A., Brown, C. & Duka, T. Attention and expectation in human predictive learning: The role of uncertainty. Quarterly Journal of Experimental Psychology 61, 1658–1668 (2008).

28 Zhuo, Y. et al. Improved green and red GRAB sensors for monitoring dopaminergic activity in vivo. Nat Methods 21, 680–691 (2024). 10.1038/s41592-023-02100-w

29 De Tommaso, M. & Turatto, M. Testing rewardLJcue attentional salience: Attainment and dynamic changes. British Journal of Psychology 113, 396–411 (2022).

30 De Tommaso, M. & Turatto, M. On the resilience of reward cues attentional salience to reward devaluation, time, incentive learning, and contingency remapping. Behavioral Neuroscience 135, 389 (2021).

31 De Tommaso, M., Mastropasqua, T. & Turatto, M. The salience of a reward cue can outlast reward devaluation. Behavioral Neuroscience 131, 226 (2017).

32 Mackintosh, N. J. A theory of attention: Variations in the associability of stimuli with reinforcement. Psychol Rev 82, 276 (1975).

33 Le Pelley, M. The hybrid modeling approach to conditioning. Computational models of conditioning, 71–107 (2010).

34 Esber, G. R. & Haselgrove, M. Reconciling the influence of predictiveness and uncertainty on stimulus salience: a model of attention in associative learning. Proceedings of the Royal Society B: Biological Sciences 278, 2553–2561 (2011).

35 Pearce, J. M. & Hall, G. A model for Pavlovian learning: variations in the effectiveness of conditioned but not of unconditioned stimuli. Psychol Rev 87, 532 (1980).

36 Sutherland, N. S. & Mackintosh, N. J. Mechanisms of animal discrimination learning. (Academic Press, 2013).

37 Pearce, J., Kaye, H. & Hall, G. Predictive accuracy and stimulus associability: Development of a model for Pavlovian learning. Quantitative analyses of behavior 3, 241–255 (1982).

38 George, D. N. & Pearce, J. M. Acquired distinctiveness is controlled by stimulus relevance not correlation with reward. Journal of Experimental Psychology: Animal Behavior Processes 25, 363 (1999).

39 Duffaud, A. M., Killcross, S. & George, D. N. Optional-shift behaviour in rats: A novel procedure for assessing attentional processes in discrimination learning. Quarterly Journal of Experimental Psychology 60, 534–542 (2007).

40 Lubow, R. E. Latent inhibition. Psychological bulletin 79, 398 (1973).

41 Lubow, R. E. Latent inhibition and conditioned attention theory. (Cambridge University Press, 1989).

42 Soltani, A. & Izquierdo, A. Adaptive learning under expected and unexpected uncertainty. Nature Reviews Neuroscience 20, 635–644 (2019).

43 Piray, P. & Daw, N. D. A model for learning based on the joint estimation of stochasticity and volatility. Nature communications 12, 6587 (2021).

44 Mah, A., Golden, C. E. & Constantinople, C. M. Dopamine transients encode reward prediction errors independent of learning rates. Cell reports 43 (2024).

45 Shen, W. et al. M4 muscarinic receptor signaling ameliorates striatal plasticity deficits in models of L-DOPA-induced dyskinesia. Neuron 88, 762–773 (2015).

46 Lowes, D. C. et al. Striatal acetylcholine enables latent-state creation during reversal learning. bioRxiv, 2026.2007. 2029.741321 (2026).

47 Apicella, P., Ravel, S., Deffains, M. & Legallet, E. The role of striatal tonically active neurons in reward prediction error signaling during instrumental task performance. Journal of Neuroscience 31, 1507–1515 (2011).

48 Sturgill, J. F. et al. Basal forebrain-derived acetylcholine encodes valence-free reinforcement prediction error. BioRxiv, 2020.2002. 2017.953141 (2020).

49 Gallo, E. F. et al. Dopamine D2 receptors modulate the cholinergic pause and inhibitory learning. Molecular psychiatry 27, 1502–1514 (2022).

